# Immunopeptidomics reveals determinants of *Mycobacterium tuberculosis* antigen presentation on MHC class I

**DOI:** 10.1101/2022.08.30.505882

**Authors:** Owen Leddy, Forest M. White, Bryan D. Bryson

**Affiliations:** Department of Biological Engineering, Massachusetts Institute of Technology, Cambridge, MA; Ragon Institute of Massachusetts General Hospital, Harvard, and MIT, Cambridge, MA; Koch Institute for Integrative Cancer Research, Cambridge, MA; Center for Precision Cancer Medicine, Cambridge, MA

## Abstract

CD8+ T cell recognition of *Mycobacterium tuberculosis* (*Mtb*)-specific peptides presented on major histocompatibility complex class I (MHC-I) contributes to immunity to tuberculosis (TB), but the principles that govern presentation of *Mtb* antigens on MHC-I are incompletely understood. In this study, mass spectrometry (MS) analysis of the MHC-I repertoire of *Mtb*-infected primary human macrophages reveals that substrates of *Mtb*’s type VII secretion systems (T7SS) are overrepresented among *Mtb*-derived peptides presented on MHC-I. Quantitative, targeted MS shows that ESX-1 activity contributes to presentation of *Mtb* antigens on MHC-I, consistent with a model in which *Mtb* T7SS substrates access a cytosolic antigen processing pathway via ESX-1-mediated phagosome permeabilization.

Chemical inhibition of proteasome activity, lysosomal acidification, or cysteine cathepsin activity did not block presentation of *Mtb* antigens on MHC-I, suggesting involvement of other proteolytic pathways or redundancy among multiple pathways. Our study identifies *Mtb* antigens presented on MHC-I that could serve as targets for TB vaccines, and reveals an important role for T7SS activity in presentation of *Mtb* antigens on MHC-I.

## Main

Tuberculosis (TB), caused by *Mycobacterium tuberculosis* (*Mtb*), is a leading cause of infectious disease mortality worldwide, causing approximately 10 million new cases of active TB disease and 1.5 million deaths per year.^1^ Currently, the only clinically licensed vaccine to prevent TB is Bacille Calmette-Guerin (BCG), which protects children against disseminated *Mtb* infection,^2^ but provides limited and highly variable protection against pulmonary TB in adults.^3^ More effective vaccines against TB are therefore needed, but identifying *Mtb* antigens capable of eliciting protective immunity remains challenging.

Multiple convergent lines of evidence from experiments in mouse and non-human primate models of TB show that CD8+ T cells can contribute to immune control of *Mtb* infection,^4–6^ but the antigenic targets of protective CD8+ T cell immunity to *Mtb* infection have not been conclusively defined. In murine models, CD8+ T cells specific for some immunodominant *Mtb* antigens poorly recognize *Mtb*-infected macrophages,^7^ implying that infected macrophages may not present all *Mtb* antigens that elicit cytokine-producing CD8+ T cell responses.^8,9^ These results suggest a need to directly identify which *Mtb* antigens are presented on MHC-I by infected phagocytes.^10^

It is currently unknown which *Mtb* antigens are presented on MHC-I in macrophages infected with virulent *Mtb*. Whereas some bacterial species are lysed following phagocytosis and expose their cell contents to antigen processing pathways,^11^ a high proportion of virulent *Mtb* remains intact and viable in macrophages,^12–14^ leading us to hypothesize that only a subset of *Mtb* proteins may be accessible for processing and presentation on MHC-I. Here, we use MS-based identification of peptides bound to MHC-I (immunopeptidomics) to directly identify *Mtb*-derived peptides presented on MHC-I in primary human macrophages infected with virulent *Mtb* H37Rv, revealing potential targets for CD8+ T cell-mediated immunity. Additionally, we use targeted MS to quantify changes in the presentation of *Mtb* peptides resulting from genetic perturbations to *Mtb* and chemical perturbations to the host cell, allowing us to probe host and bacterial determinants of antigen presentation on MHC-I in *Mtb* infection.

To identify *Mtb* antigens presented on MHC-I, we infected primary human monocyte-derived macrophages with *Mtb* H37Rv, isolated MHC-I by immunoprecipitation 72 hours post-infection, purified the associated peptides, and identified peptides by liquid chromatography coupled to tandem mass spectrometry (LC-MS/MS). For an initial set of 3 analyses, we adapted a previously described protocol^15^ (Protocol 1 − see Methods) for use in a biosafety level 3 (BL3) setting. We developed a further optimized protocol (Protocol 2 − see Methods) that we used for 3 additional analyses (Figure 1a). The class I human leukocyte antigen (HLA) genotypes of the 6 donors analyzed were largely distinct from one another (Supplementary Table 1), enabling us to identify peptides associated with a variety of HLA alleles.

**Figure 1.**
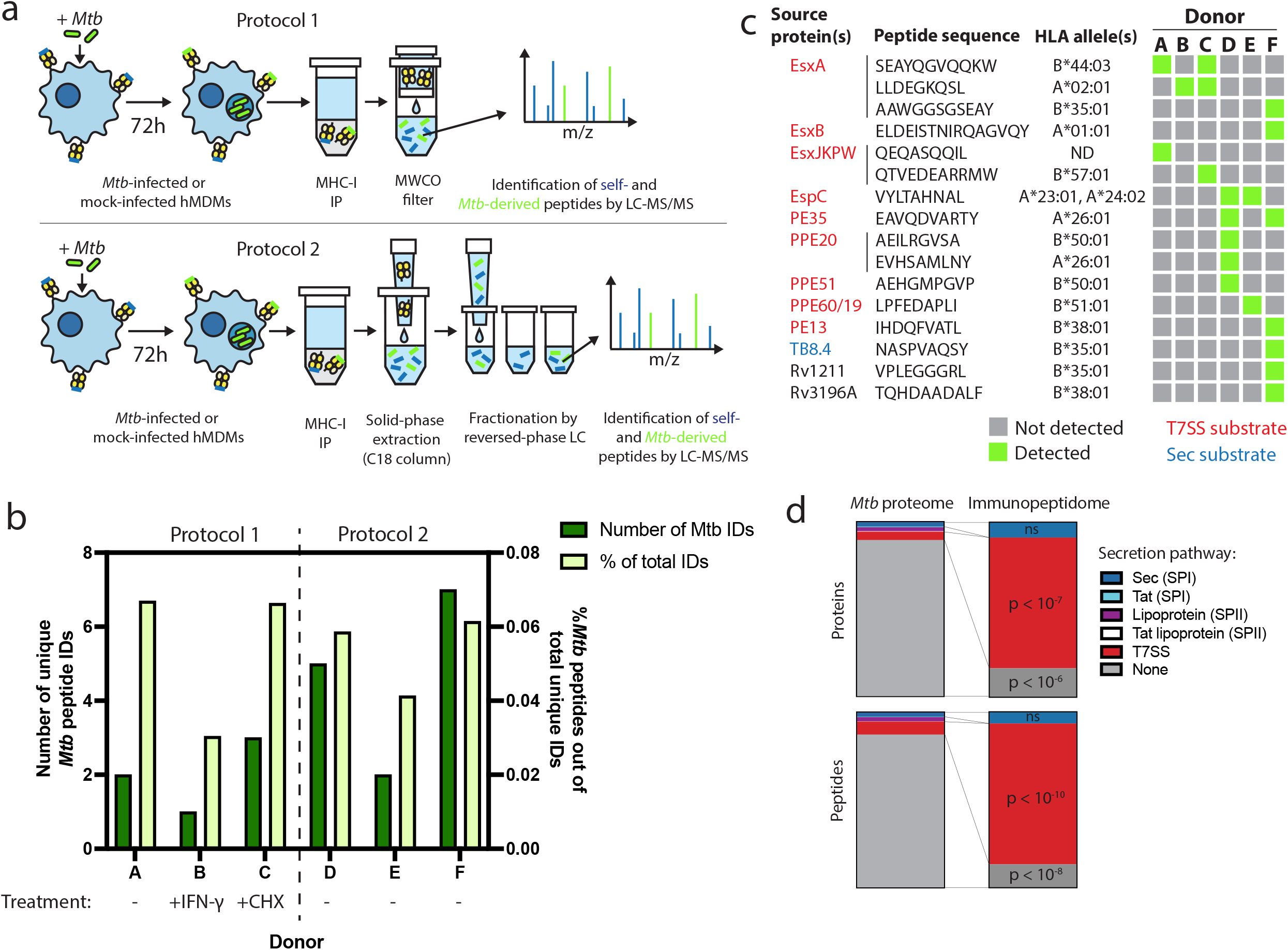
The MHC-I immunopeptidome of *Mtb*-infected human macrophages is enriched for T7SS substrates. a) Schematic representation of two immunopeptidomics workflows used to profile the MHC-I repertoire of *Mtb*-infected primary human macrophages. IP: immunoprecipitation. MWCO: molecular weight cutoff. LC-MS/MS: liquid chromatography coupled to tandem mass spectrometry. b) Absolute and relative number of *Mtb*-derived MHC-I peptides identified for each donor. Macrophages from donor B were pre-treated for 24 hours with 10 ng/mL IFN-γ, and macrophages from donor C were treated with 0.5 µg/mL cycloheximide (CHX) for the final 6 hours of infection. c) Sequences, source proteins, associated HLA alleles, and donors for each validated *Mtb*-derived MHC-I peptide. d) Enrichment analysis of *Mtb* peptides presented on MHC-I and their source proteins, categorized by protein secretion pathway using SignalP 6.0^54^ and a curated set of known or strongly suspected T7SS substrates. p-values for enrichment analyses of proteins and peptides were determined using the binomial test and the hypergeometric test respectively (see Methods).

We identified thousands of peptides for each primary cell donor, with a length distribution typical of MHC-I peptides (Supplementary Fig. 1 a) and retention times that correlated well with hydrophobicity (Supplementary Fig. 1 b). Unsupervised clustering of identified peptides using GibbsCluster 2.0^16^ revealed groups corresponding to the known peptide sequence binding motifs of class I HLA alleles expressed by each donor (Supplementary Fig. 2), confirming the specificity of our pulldowns. *Mtb-*derived MHC-I peptides detected in each pulldown were predicted to bind at least one class I HLA allele expressed by the donor (Supplementary Fig. 3). *Mtb* peptides made up less than 0.1% of MHC-I peptides identified, and this proportion was not discernibly increased by pre-treating macrophages with IFN-γ or treating with cycloheximide to inhibit host protein synthesis (Figure 1b).

Putative *Mtb* peptides that passed manual inspection of MS/MS spectra and extracted ion chromatograms (see Methods) were further validated using internal standard parallel reaction monitoring (IS-PRM, also known as SureQuant)^17,18^ (Supplementary Fig. 4). Putative *Mtb* peptides that had MS/MS spectra that closely matched that of a synthetic stable isotope-labeled (SIL) standard, co-eluted with the SIL standard, and were not detected in mock-infected control samples by SureQuant were considered correctly identified, authentic *Mtb* peptides (Supplementary Fig. 5). 77.85% of MASCOT identifications of putative *Mtb* peptides were rejected after manual data inspection, and of the remaining candidates a further 51.51% were rejected following analysis by SureQuant (Supplementary Table 2), highlighting the need for rigorous validation using best practices^19^ when identifying pathogen-derived peptides among an MHC repertoire dominated by host-derived peptides.

Of the 16 *Mtb*-derived MHC-I peptides we identified, 13 (81.25%) derived from proteins secreted via type VII secretion systems (T7SS) (Fig. 1c). The *Mtb* genome encodes five of these protein export machines (designated ESX-1, 2, 3, 4, and 5),^20^ and we identified MHC-I peptides derived from proteins known to be secreted by three of these systems (ESX-1,^21,22^ ESX-3,^23^ and ESX-5^24–26^). For several T7SS substrates, we identified multiple peptides from the same protein, and/or identified the same peptide across multiple donors. These antigens included EsxA, PPE20, EspC, and PE35, as well as sequences conserved among the four nearly identical members of the EsxJ family of proteins (EsxJ, EsxK, EsxP, and EsxW − referred to here as EsxJKPW). T7SS substrates were significantly overrepresented in the MHC-I repertoire relative to the whole *Mtb* proteome, both at the peptide level (p < 10^−10^; binomial test with Bonferroni correction) and at the protein level (p < 10^−7^; hypergeometric test with Bonferroni correction) (Fig. 1d). Proteins without identifiable secretion signals were significantly underrepresented among *Mtb-* derived MHC-I peptides (p < 10^−8^; binomial test with Bonferroni correction) and source proteins (p < 10^−6^; hypergeometric test with Bonferroni correction). While some of the *Mtb* antigens we identified are highly abundant compared to the rest of the *Mtb* proteome (e.g., EsxB), the abundances of other T7SS substrates are near the mean (Supplementary Figure 1c),^27^ suggesting that the overrepresentation of T7SS substrates is not solely due to greater abundance relative to the rest of the *Mtb* proteome. These results suggest that T7SS substrates may preferentially gain access to MHC-I antigen processing pathways.

To nominate possible mechanisms by which *Mtb* antigens might be processed and loaded onto MHC-I, we used confocal microscopy to examine the intracellular fate of *Mtb* in primary human macrophages. We hypothesized that *Mtb* antigens could either (1) gain access to cytosolic antigen processing pathways via permeabilization of the phagosome membrane, or (2) could be processed by endolysosomal proteases and loaded onto MHC-I in *Mtb*-containing compartments (Fig. 2 a). In a subset of macrophages at both early and late timepoints, a subset of *Mtb* co-localized with Galectin-3, a marker of phagosomal membrane permeabilization^28,29^ (Fig. 2 b, c; Supplementary Fig. 6 a). A subset of *Mtb* also co-localized with P62, an autophagy adaptor protein that recognizes cytosol-exposed bacteria^30^ (Fig. 2 b, c; Supplementary Fig. 6 b). Since bacteria become ubiquitinated upon exposure to the cytosol^28^ and recruitment of autophagy adaptor proteins to the *Mtb* phagosome has previously been associated with damage to the phagosomal membrane,^28^ these results suggest that *Mtb* gains access to the host cytosol in primary human macrophages, as has previously been shown in murine macrophages^31^ and in cell lines.^32^ A subset of *Mtb*-containing phagosomes co-localized with the late endosome and lysosome marker LAMP-1 (Fig. 2 b, c; Supplementary Fig. 6 c), suggesting access to endolysosomal proteases, but *Mtb*-containing phagosomes did not co-localize with MHC-I itself (Fig. 2 b, c; Supplementary Fig. 6 d), suggesting that *Mtb* antigens were unlikely to be loaded onto MHC-I in the *Mtb*-containing phagosome. These results suggested that access to the host cell cytosol represented a likely route for processing and presentation of *Mtb* antigens. An *Mtb* strain deficient in the activity of the ESX-1 secretion system (*eccCa1:Tn*) showed reduced co-localization with Galectin-3 and P62 (Fig. 2 d, e, f), consistent with the known role of ESX-1 activity in enabling *Mtb* to damage the phagosomal membrane and gain access to the host cytosol.^33^

**Figure 2.**
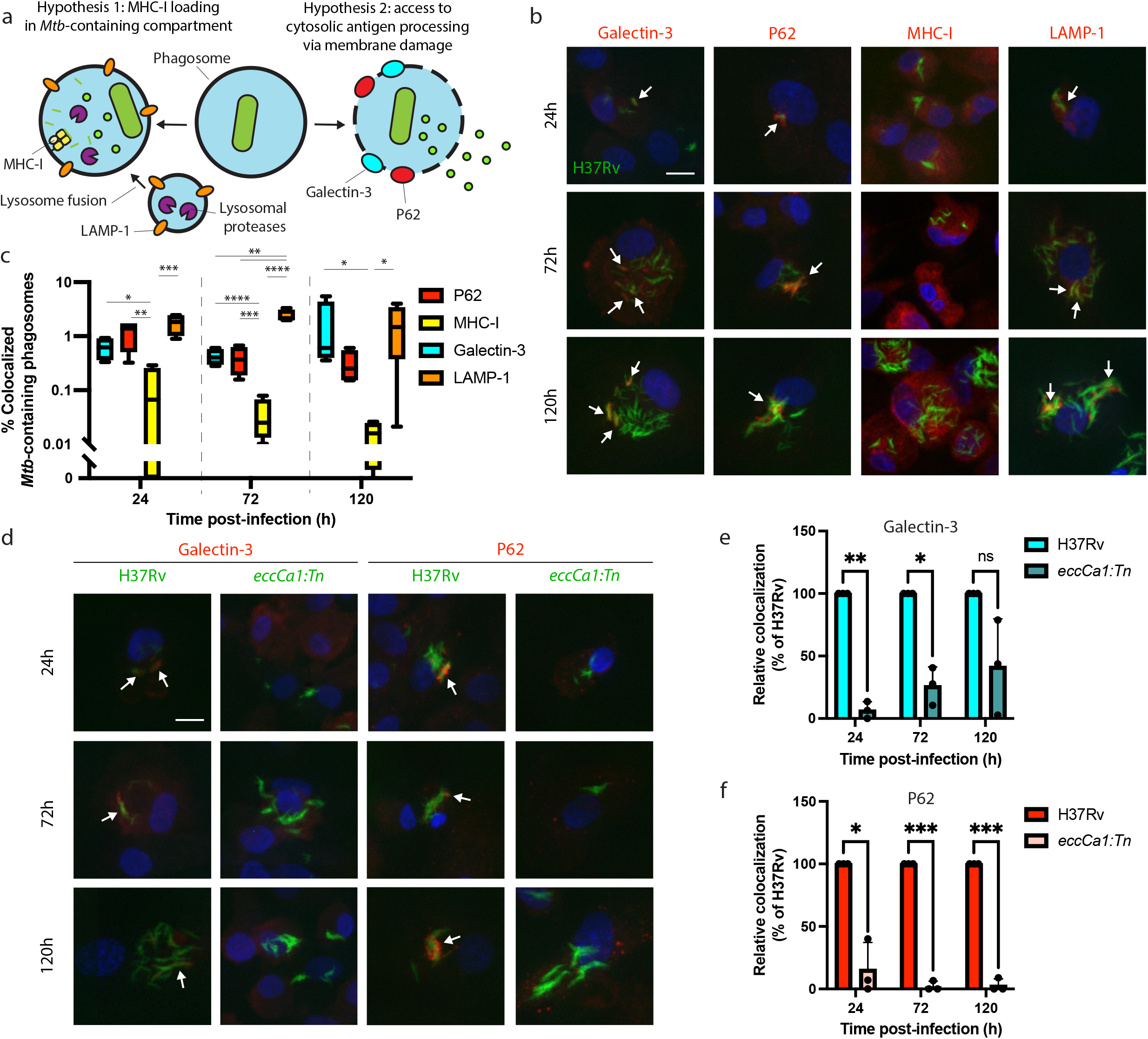
*Mtb* co-localizes with markers of phagosome membrane damage in an ESX-1-dependent manner and does not co-localize with MHC-I. Primary human macrophages were infected with GFP-expressing wild-type *Mtb*, fixed at 24, 72, or 120 hours post-infection, stained by immunofluorescence (IF), and imaged by spinning-disk confocal microscopy. a) Schematic showing markers associated with each of two possible pathways of *Mtb* antigen processing and presentation. b) Representative images of *Mtb-*infected macrophages stained for Galectin-3, P62, MHC-I, or LAMP-1. Scale bar indicates 10 µm. White arrows indicate *Mtb*-containing phagosomes co-localizing with each marker. c) Automated quantification (see Methods) of the proportion of co-localized *Mtb*-containing phagosomes for each marker for n=4 donors (* p < 0.05, ** p < 0.01, *** p < 0.001, **** p < 0.0001; one-way ANOVA with Tukey’s multiple comparisons test). d) Representative image of macrophages infected with wild-type (H37Rv) or ESX-1-deficient (*eccCa1:Tn*) *Mtb* stained for Galectin-3 or P62. Scale bar indicates 10 µm. White arrows indicate *Mtb*-containing phagosomes co-localizing with each marker. e-f) Automated quantification of the relative proportion of GFP+ objects co-localized with IF staining for Galectin-3 (e) or P62 (f) as a function of time post-infection for n = 3 donors, normalized to wild-type (H37Rv) (* p < 0.05, ** p < 0.01, *** p < 0.001; paired t-test).

Our microscopy results led us to hypothesize that ESX-1-mediated phagosomal membrane damage might be required for *Mtb* antigens to access MHC-I antigen processing pathways (Fig. 3 a). If this were the case, ESX-1 activity would be required for presentation not only of peptides derived from ESX-1 substrates (e.g., EsxA_28-36_ − sequence LLDEGKQSL), but also peptides derived from substrates of other T7SSs (e.g., EsxJKPW_24-34_ − sequence QTVEDEARRMW − which is derived from ESX-5 substrates that do not require ESX-1 for secretion)^34^.To test this hypothesis, we turned to quantitative targeted MS to quantify changes in the presentation of EsxA_28-36_ and EsxJKPW_24-34_ across multiple experimental conditions (Fig. 3 b). We used primary macrophages from donors expressing HLA-A*02:01 and HLA-B*57:01 for these experiments to ensure presentation of the target peptides. We spiked pre-formed heavy isotope labeled peptide-MHC complexes (hipMHCs) into the lysates prior to immunoprecipitation to provide an internal standard that can be used to normalize out technical variation, which improves the accuracy of label-free quantification.^15^

**Figure 3.**
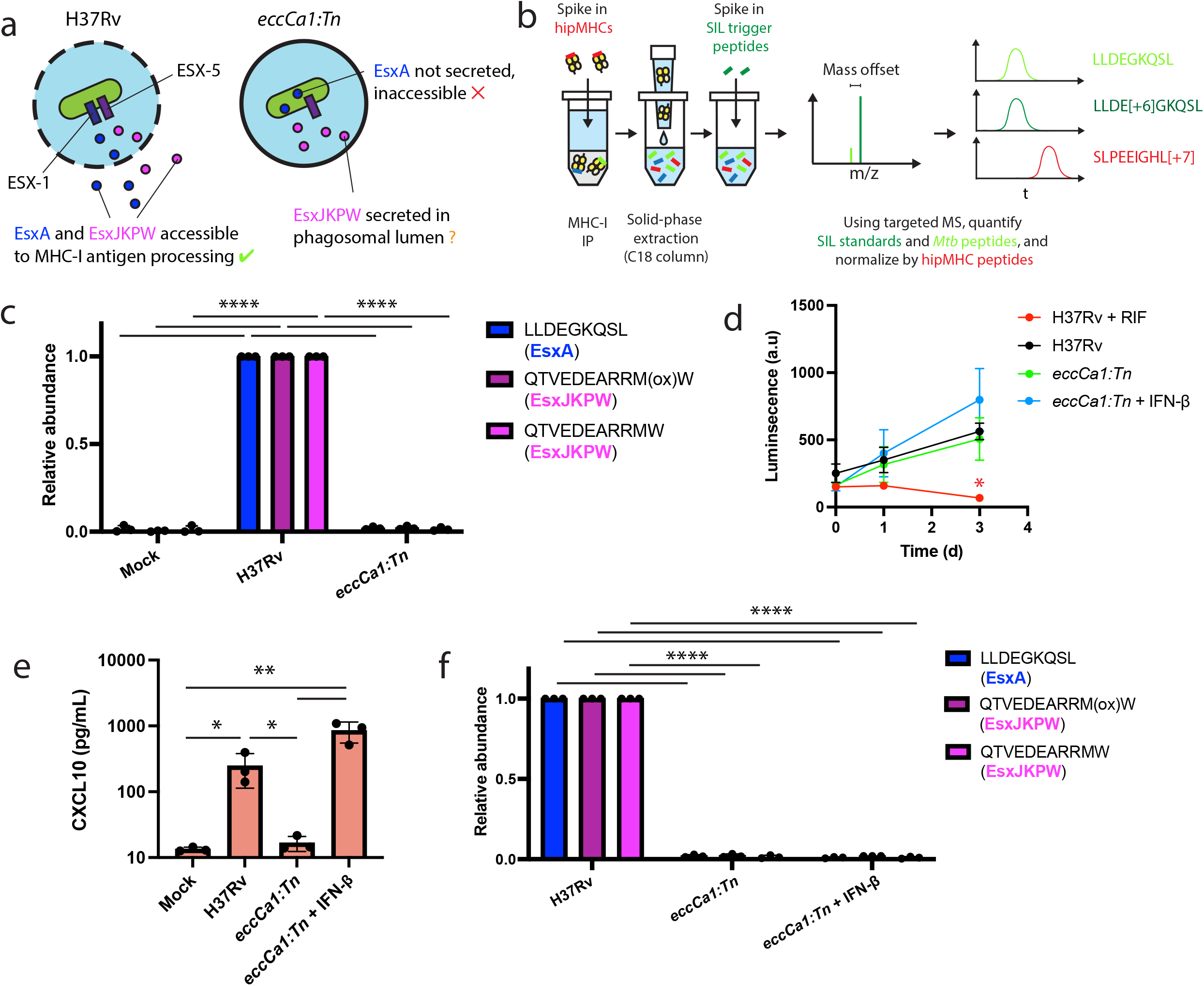
ESX-1 activity is required for presentation of EsxA_28-36_ and EsxJKPW_24-34_ on MHC-I, independently of type I interferon signaling. a) Schematic representation of the localization of EsxA and EsxJKPW in macrophages infected with wild-type *Mtb* H37Rv or the ESX-1-deficient *eccCa1:Tn* transposon mutant. b) Schematic showing our workflow for targeted detection and quantification of *Mtb*-derived MHC-I peptides by SureQuant, using heavy isotope labeled peptide-MHC complexes (hipMHCs) as internal standards. SIL: stable isotope labeled. c) Relative quantification of EsxA_28-36_ and EsxJKPW_24-34_ by SureQuant in macrophages infected with no *Mtb* (mock), wild-type *Mtb* H37Rv, or *eccCa1:Tn* for n = 3 donors (all HLA-A*02:01+, HLA-B*57:01+). As oxidation of methionine is common during sample handling, both the oxidized and non-oxidized form of EsxJKPW_24-34_ were quantified. d) Luminescence as a function of time measured for macrophages infected with luciferase-expressing *Mtb*, in a wild-type H37Rv or *eccCa1:Tn* background, with or without the addition of 10 ng/mL IFN-β in the culture media. Addition of 25 ug/mL rifampicin (RIF) to the culture media was used as a control showing reduced luminescence with bacterial death. Data points and error bars represent the mean and standard deviation of n = 3 donors, each of which represents the mean of three technical replicates. (* p < 0.05, one-way ANOVA with Dunnett’s multiple comparisons test, relative to H37Rv as the reference condition). e) CXCL10 concentration in the culture media 72 hours post-infection quantified by ELISA. Data points each represent the mean of three technical replicates for a given donor. (* p < 0.05, ** p < 0.01, one-way ANOVA with Tukey’s multiple comparisons test on log-transformed concentrations.) f) Relative quantification of EsxA_28-36_ and EsxJKPW_24-34_ by SureQuant in macrophages infected with no *Mtb* (mock), wild-type *Mtb* H37Rv, or *eccCa1:Tn* for n = 3 donors (all HLA-A*02:01+, HLA-B*57:01+). (**** p < 0.001, one-way ANOVA with Tukey’s multiple comparisons test.)

Targeted MS analysis revealed that macrophages infected with ESX-1-deficient *Mtb* (*eccCa1:Tn*) did not present either EsxA_28-36_ or EsxJKPW_24-34_ on MHC-I (Fig. 3c). We assessed alternative explanations for this difference in presentation of EsxA_28-36_ and EsxJKPW_24-34_ and observed no difference in *Mtb* outgrowth (Fig. 3d), the proportion of cells infected (Supplementary Fig. 7 a, b), or total MHC-I surface expression (Supplementary Fig. 7 c, d) between macrophages infected with wild-type or ESX-1-deficient *Mtb*. These results show that ESX-1 activity is required for presentation of EsxA_28-36_ and EsxJKPW_24-34_ on MHC-I in macrophages.

To determine whether the absence of EsxA_28-36_ and EsxJKPW_24-34_ in the MHC-I repertoire of macrophages infected with ESX-1-deficient *Mtb* could be attributed to a loss of type I interferon signaling,^35^ we added exogenous IFN-β to macrophages infected with ESX-1-deficient *Mtb* and again quantified presentation of EsxA_28-36_ and EsxJKPW_24-34_ by SureQuant. Addition of exogenous IFN-β restored a type I interferon response as measured by production of CXCL10 (Fig. 3e), but did not rescue presentation of EsxA_28-36_ or EsxJKPW_24-34_ on MHC-I (Fig. 3f). These results show that presentation of EsxA_28-36_ or EsxJKPW_24-34_ on MHC-I is dependent on ESX-1 activity but independent of downstream type I interferon signaling, consistent with a model in which ESX-1-mediated phagosomal damage enables *Mtb* antigens to access MHC-I antigen processing pathways.

Many MHC-I peptides are proteolytically processed by the proteasome,^36^ while others are processed by endosomal or lysosomal proteases.^37,38^ To determine whether these mechanisms contribute to presentation of *Mtb* peptides on MHC-I, we treated HLA-A*02:01+, HLA-B*57:01+ macrophages with inhibitors of proteasome activity (MG-132), cysteine cathepsin activity (E64d), and lysosomal acidification (bafilomycin) (Fig. 4a). We quantified presentation of EsxA_28-36_ and EsxJKPW_24-34_ on MHC-I along with five host-derived HLA-A*02:01-binding peptides identified in previous studies^39^ that could be reliably detected in the MHC-I repertoire of macrophages. We began drug treatment of the macrophages prior to infection with *Mtb* and limited the duration of infection to 24h so that the cells could be treated with drug for the full duration of the infection without excessive cytotoxicity (Fig. 4b). Treatment with MG-132 inhibited proteasome activity, as measured by accumulation of proteins modified with K48-linked polyubiquitin (Supplementary Fig. 8 a, b). Treatment with E64d inhibited cathepsin B activity, as measured by a fluorometric assay (Supplementary Fig. 8 c). Treatment with bafilomycin inhibited lysosomal acidification, as measured by lysotracker staining (Supplementary fig 8 d, e). All three drugs exhibited minimal cytotoxicity in macrophages at the doses used in our immunopeptidomic experiments (Supplementary Fig. 9), and did not inhibit phagocytosis of *Mtb* or bacterial outgrowth (Supplementary Fig. 10).

**Figure 4.**
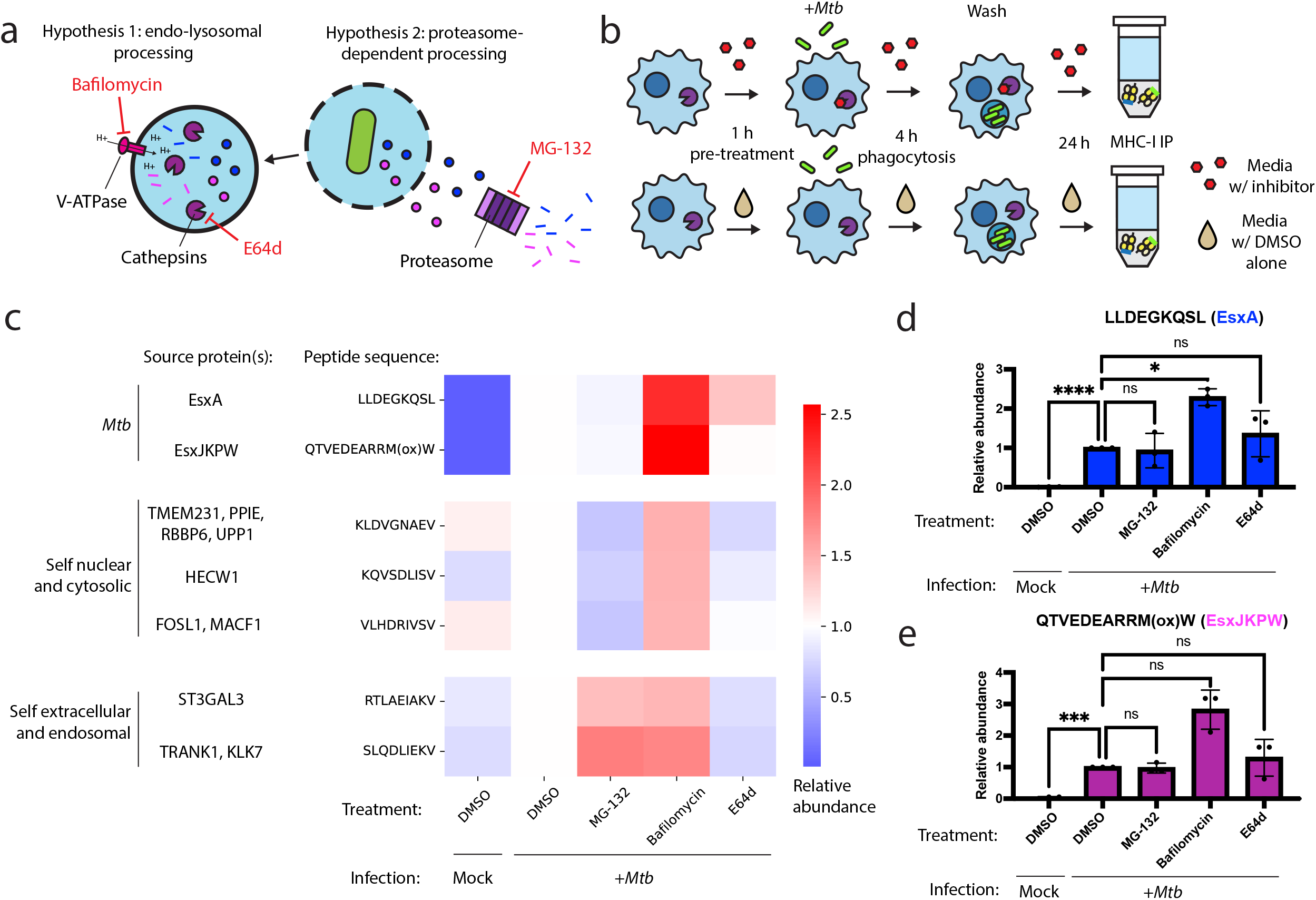
Inhibition of conventional antigen processing proteolytic pathways does not impair presentation of EsxA_28-36_ and EsxJKPW_24-34_ on MHC-I. a) Schematic representation of proteolytic pathways inhibited by the proteasome inhibitor MG-132, the V-type ATPase inhibitor bafilomycin, and the cysteine cathepsin inhibitor E64d. b) Schematic showing the timing of drug treatment and *Mtb* infection for targeted MS experiments. c) Heatmap showing relative abundance of self and *Mtb*-derived MHC-I peptides determined by SureQuant in mock-infected macrophages or *Mtb*-infected macrophages treated with MG-132, bafilomycin, E64d, or DMSO-only control. Colors represent the mean fold change relative to the DMSO-treated, *Mtb*-infected condition for n = 3 donors (all HLA-A*02:01+, HLA-B*57:01+). d-e) Relative abundance of EsxA_**28-36**_ (d) and EsxJKPW_24-34_ (e) determined by SureQuant in mock-infected macrophages or *Mtb*-infected macrophages treated with MG-132, bafilomycin, E64d, or DMSO-only control for n = 3 donors (all HLA-A*02:01+, HLA-B*57:01+).

Treatment with MG-132 reduced presentation of MHC-I peptides derived from cytosolic or nuclear host proteins, but not peptides derived from *Mtb* proteins or secreted or endomembrane compartment localized host proteins (Fig. 4c). Treatment with E64d decreased presentation of peptides derived from endosomal or secreted proteins and had a range of effects on peptides derived from cytosolic or nuclear proteins, but did not decrease presentation of *Mtb* peptides (Fig. 4c). Treatment with bafilomycin broadly increased presentation of target MHC-I peptides (Fig. 4c). This increase was statistically significant for EsxA_28-36_ (Fig. 4d; p < 0.05, one-way ANOVA with Dunnett’s multiple comparisons test), though not for EsxJKPW_24-34_ (Fig. 4e). These results suggest that processing of *Mtb* peptides for presentation on MHC-I relies on antigen processing proteases other than the proteasome, cysteine cathepsins, or other acidification-dependent lysosomal proteases, and/or that multiple redundant pathways contribute to processing of *Mtb* peptides for presentation on MHC-I.

## Discussion

Our analysis of the MHC-I repertoire of primary human macrophages infected with *Mtb* revealed that T7SS substrates are a prominent source of *Mtb* peptides presented on MHC-I. Our findings contrast with those of previous studies of the immunopeptidome of macrophage cell lines infected with BCG^40^ and the avirulent strain *Mtb* H37Ra,^41^ in which the mycobacterial peptides identified included twin arginine translocation (Tat) pathway substates, Sec pathway secretion substrates, membrane-associated proteins, and cytosolic proteins, with no apparent enrichment for T7SS substrates. Given that we showed ESX-1 activity contributes to presentation of *Mtb* antigens in infection with virulent *Mtb* H37Rv, the fact that BCG and *Mtb* H37Ra are both deficient in ESX-1 activity^21,42^ may in part explain this difference in the mycobacterial peptides presented on MHC-I.

Our findings contrast with previous work arguing that ESX-1 activity is dispensable for presentation of *Mtb* antigens on MHC-I. ESX-1 activity is not required for human monocyte-derived dendritic cells to present an epitope derived from TB8.4 (a Sec pathway substrate) to cognate T cells when infected with *Mtb*,^43^ or for priming of CD8+ T cell responses specific for non-ESX-1-secreted antigens *in vivo* in mice.^44^ While these prior results suggest that the requirements for presentation of *Mtb* antigens on MHC-I may vary among *Mtb* antigens and/or vary among antigen presenting cell types, our results show that ESX-1 activity is essential for the presentation of certain *Mtb* antigens in infected human macrophages. Whereas in axenic culture, secretion of ESX-5 substrates is independent of ESX-1 function,^34,45^ the fact that presentation of peptides derived from an ESX-5 substrate on MHC-I requires ESX-1 activity suggests that the localization of secreted *Mtb* proteins within a host cell may depend on the activity of multiple T7SSs. While our results are consistent with a model in which ESX-1-mediated phagosomal membrane damage enables *Mtb* antigens to access cytosolic antigen processing and presentation pathways, further evidence will be needed to establish whether phagosomal membrane damage alone is sufficient to enable presentation of *Mtb* antigens on MHC-I in the absence of other ESX-1 functions.

In addition to T7SS substrates, we also identified peptides derived from a Sec pathway substrate (TB8.4), and from uncharacterized hypothetical proteins (Rv1211, Rv3196A). Rv1211 and Rv3196A have both previously been detected by MS in culture filtrates of *Mtb*,^46^ suggesting they could be secreted despite lacking readily identifiable secretion signals. TB8.4, Rv1211, and Rv3196A are all low molecular weight proteins comparable in size to the Esx family of proteins, which we speculate could facilitate translocation across a permeabilized phagosomal membrane. Further investigation of the localization of these proteins may help us better understand how and to what extent small secreted *Mtb* proteins other than known T7SS substrates can contribute to the MHC-I repertoire.

Some of the *Mtb* MHC-I peptides we identified derive from proteins known or suspected to be associated with the mycobacterial outer membrane. These include PPE51,^47^ PPE20,^48^ PPE60,^49^ EspC,^50^ and PE13 (the co-transcribed binding partner^51^ of the putative outer membrane protein PPE18^52^). While some of these proteins can also be found in soluble form in culture filtrates,^23^ others may be stably associated with the mycobacterial outer membrane.^47^ The mechanism by which antigens associated with the outer membrane are processed and presented could differ from that of soluble secreted proteins.

Prior work shows that at least some of the MHC-I antigens we identified as potential vaccine targets are immunogenic in humans. Lewinsohn et al.^8^ showed that in humans with prior exposure to *Mtb*, the loci that encode EsxA, EsxB, PPE51, and EsxJKPW are sources of immunodominant CD8+ T cell antigens in *Mtb*-exposed humans, while other antigens identified in our study elicited intermediate responses (PE13, PE35, PPE60, PPE19), and one elicited no detectable response (PPE20). In a separate study, CD8+ T cells (as well as CD4+ T cells) recognized multiple epitopes derived from EspC.^22^ Antigens like PPE20 that did not elicit measurable CD8+ T cell responses in the context of natural infection should not necessarily be ruled out as vaccine targets, as a lack of CD8+ T cell recognition in the setting of natural infection does not necessarily imply that these antigens could not be immunogenic.

Our results identify several *Mtb* proteins that are accessible for presentation on MHC-I in *Mtb*-infected human macrophages, suggesting that these could serve as targets for subunit vaccines designed to elicit CD8+ T cell-mediated protection against TB. By showing that T7SS substrates are overrepresented in the MHC-I repertoire of macrophages and revealing mechanistic determinants of *Mtb* antigen processing, our results have the potential to guide future screens to identify additional vaccine targets. Our findings advance the field’s understanding of antigen presentation in TB and suggest novel potential targets for TB vaccine development.

## Methods

### Human cell isolation, differentiation, and culture

Deidentified buffy coats were obtained from Massachusetts General Hospital. Samples are acquired and provided to research groups with no identifying information. HLA-A*02:01+, HLA-B*57:01+ leukapheresis samples (leukopaks)were obtained from StemCell. PBMCs were isolated by density-based centrifugation using Ficoll (GE Healthcare). CD14+ monocytes were isolated from PBMCs using a CD14 positive-selection kit (Stemcell). Isolated monocytes were differentiated in R10 media [RPMI 1640 without phenol red (Gibco) supplemented with 10% heat-inactivated FBS (Gibco), 1% HEPES (Corning), 1% L-glutamine (Sigma)] supplemented with 25 ng/mL M-CSF (Biolegend, 572902). Monocytes were cultured on appropriate ultra-low-attachment plates or flasks (Corning) for 6 days.

For ELISA assays, Cathepsin B activity assays, and growth curves (see below), after 6 days macrophages were detached using a detachment buffer of calcium-free PBS and 2 mM ethylenediaminetetraacetic acid (EDTA), pelleted, and recounted. Macrophages were plated in tissue culture treated 96-well plates at a density of 50,000 cells per well in R10 media and allowed to re-adhere overnight prior to infection. For microscopy (see below) cells were similarly detached and were replated on 12-well chamber slides (Ibidi) at a density of 100,000 cells per well in R10 media and allowed to re-adhere overnight prior to infection. For all other experiments, macrophage differentiation media was replaced with fresh R10 overnight prior to infection, without re-plating the macrophages.

### HLA genotyping

Genomic DNA was extracted from 5 × 10^6^ PBMCs using a Qiagen DNeasy kit. HLA typing was performed using a targeted next generation sequencing (NGS) method. Briefly, locus-specific primers were used to amplify a total of 26 polymorphic exons of HLA-A & B (exons 1 to 4), C (exons 1 to 5), E (exon 3), DPA1 (exon 2), DPB1 (exons 2 to 4), DQA1 (exon 1 to 3), DQB1 (exons 2 & 3), DRB1 (exons 2 & 3), and DRB3/4/5 (exon 2) genes with Fluidigm Access Array system (Fluidigm Corporation, South San Francisco, CA 94080 USA). The 26 Fluidigm PCR amplicons were harvested from Fluidigm Access Allay IFC and pooled. Quality and quantity were checked using a Caliper LabChip GX Touch HT Nucleic Acid Analyzer (PerkinElmer, Waltham, MA 02452 USA). The PCR product library was quantitated and subjected to sequencing on an Illumina MiSeq sequencer (Illumina, San Diego, CA 92122 USA). HLA alleles and genotypes were called using the Omixon HLA Explore (version 2.0.0) software (Omixon Biocomputing Ltd., Budapest, Hungary).

### *Mtb* culture

Mycobacterium tuberculosis (Mtb) H37Rv was grown in Difco Middlebrook 7H9 media supplemented with 10% OADC, 0.2% glycerol, and 0.05% Tween-80 to mid-log phase. Strains expressing GFP were grown in media supplemented with 50 μg/mL hygromycin B (Sigma-Aldrich), and strains expressing *luxABCDE* were grown in media supplemented with 20 μg/mL Zeocin (Thermo).

### *Mtb* infection of macrophages

The *Mtb* culture was pelleted by centrifugation, washed once with PBS, resuspended in R10 media and filtered through a 5 μm syringe filter to obtain a single-cell suspension. Macrophages were infected at the indicated multiplicity of infection (MOI) for 4 h and then washed with PBS to remove extracellular *Mtb*. Infected macrophages were cultured in R10 media for the remainder of the experiment.

### Drug treatment of macrophages

Where indicated, macrophages were pre-treated with R10 media containing 20 μM E64d (Cayman), 0.5 μM MG-132 (Cayman), or 10 nM bafilomycin (InvivoGen), or DMSO alone (or other concentration of drug where indicated). The culture media was supplemented with an equal concentration of drug during infection with *Mtb*, and after washing off extracellular *Mtb*. Where indicated, the culture media was supplemented with 10 ng/mL of IFN-β after infection with *Mtb*.

### hipMHC internal standard preparation

UV-mediated peptide exchange and quantification of hipMHC complexes by ELISA were performed as previously described.^15^ Amidated peptides with sequences AL[+7]ADGVQKV-NH_2_, ALNEQIARL[+7]-NH_2_, and SLPEEIGHL[+7]-NH_2_ were used to make hipMHC standards.

### MHC-I immunoprecipitation (IP)

50 million macrophages per condition (data-dependent analysis) or 10 million macrophages per condition (quantitative SureQuant analysis) were cultured as described above in ultra-low-attachment T75 flasks (Corning). macrophages were infected at MOI 2.5 as described above, or mock-infected with media containing no *Mtb*. Where indicated for specific DDA MS experiments (donor B), the culture media was supplemented with 10 ng/mL of IFN-γ for 24 hours prior to infection with *Mtb*. Where indicated for specific DDA MS experiments (donor C), the culture media was supplemented with 0.5 µg/mL of cycloheximide for the final 6 hours of culture prior to MHC-I isolation. 72 hours post-infection, cells were detached using PBS supplemented with 2 mM EDTA, washed with PBS, and lysed in 1 mL (DDA analysis) or 0.5 mL (SureQuant analysis) of MHC lysis buffer [20 mM Tris, 150 mM sodium chloride, pH 8.0, supplemented with 1% CHAPS, 1x HALT protease and phosphatase inhibitor cocktail (Pierce), and 0.2 mM phenylmethylsulfonyl fluoride (Sigma-Aldrich)]. Lysate was sonicated using a Q500 ultrasonic bath sonicator (Qsonica) in five 30-second pulses at an amplitude of 60%, cleared by centrifugation at 16,000 x g, and sterile filtered twice using 0.2 μm filter cartridges (Pall NanoSep). For quantitative SureQuant analyses, the protein concentration of the lysates was normalized by BCA assay (Pierce), and 100 fmol of each hipMHC standard was added to each sample. Lysates were then added to protein A sepharose beads pre-conjugated to pan-MHC-I antibody (clone W6/32), prepared as previously described,^15^ and incubated rotating at 4°C overnight (12-14 hours). Beads were then washed and peptide-MHC complexes eluted as previously described.^15^

### Purification of MHC-I-associated peptides

#### Protocol 1

MHC-I-associated peptides were purified using 10 kDa molecular weight cutoff filters (Pall NanoSep) as previously described,^15^ snap-frozen in liquid nitrogen, and lyophilized.

#### Protocol 2

C18 SpinTips (Pierce) were washed with 0.1% trifluoroacetic acid, activated with 90% acetonitrile supplemented with 0.1% formic acid, and washed with 0.1% formic acid. Eluate from MHC-I IPs was applied to the column by centrifugation. The column was washed with 0.1% formic acid, and peptides were eluted by applying elution solvent (30% acetonitrile supplemented with 0.1% formic acid) by centrifugation twice. Eluates were snap-frozen in liquid nitrogen and lyophilized. Lyophilized peptides were resuspended in solvent A2 [10 mM triethyl ammonium bicarbonate (TEAB), pH 8.0] and loaded onto a fractionation column (a 200 μm inner diameter fused silica capillary packed in-house with 10 cm of 10 μM C18 beads). Peptides were fractionated using an Agilent 1100 series liquid chromatograph using buffers A2 (10 mM TEAB, pH 8.0) and B2 (99% acetonitrile, 10 mM TEAB, pH 8.0). The fractionation column was washed with solvent A2, and peptides were separated using a gradient of 1-5% solvent B2 over 5 minutes, 5-40% over 60 minutes, 40-70% over 10 minutes, hold for 9 minutes, and 70%-1% over 1 minute. 90-second fractions were collected, concatenated into 12 tubes, over 90 minutes in total. Fractions were flash-frozen in liquid nitrogen and lyophilized.

### DDA MS analyses

MHC-I peptide samples were resuspended in 0.1% formic acid. 25% of the sample was refrozen and reserved for later SureQuant validation analyses, while 75% of the sample was used for DDA analysis. For all MS analyses, samples were analyzed using an Orbitrap Exploris 480 mass spectrometer (Thermo Scientific) coupled with an UltiMate 3000 RSLC Nano LC system (Dionex), Nanospray Flex ion source (Thermo Scientific), and column oven heater (Sonation). The MHC peptide sample was loaded onto a fused silica capillary chromatography column with an integrated electrospray tip (∼1 μm orifice) prepared and packed in-house with 10 cm of 1.9 μm C18 beads (ReproSil-Pur).

Standard mass spectrometry parameters were as follows: spray voltage, 2.0 kV; no sheath or auxiliary gas flow; ion transfer tube temperature, 275 °C. The Orbitrap Exploris 480 mass spectrometer was operated in data dependent acquisition (DDA) mode. Peptides were eluted using a gradient of 6-25% buffer B (70% Acetonitrile, 0.1% formic acid) over 75 minutes, 25-45% over 5 minutes, 45-100% over 5 minutes, hold for 1 minutes, and 100% to 3% over 2 minutes. Full scan mass spectra (350-1200 m/z, 60,000 resolution) were detected in the orbitrap analyzer after accumulation of 3 × 10^6^ ions (normalized AGC target of 300%) or 25 ms. For every full scan, MS/MS scans were collected during a 3 second cycle time. Ions were isolated (0.4 m/z isolation width) using the standard AGC target and automatic determination of maximum injection time, fragmented by HCD with 30% CE, and scanned at a resolution of 45,000. Charge states < 2 and > 4 were excluded, and precursors were excluded from selection for 30 seconds if fragmented n=2 times within a 20 second window.

### DDA data search and manual inspection

All mass spectra were analyzed with Proteome Discoverer (PD, version 2.5) and searched using Mascot (version 2.4) against a custom database comprising the Uniprot human proteome (UP000005640) and the Uniprot *Mycobacterium tuberculosis* H37Rv proteome (UP000001584). No enzyme was used, and variable modifications included oxidized methionine for all analyses. Peptide-spectrum matches from all analyses were filtered with the following criteria: search engine rank = 1, isolation interference ≤ 30%, length between 8 and 16 amino acids, ions score ≥ 15, and percolator q-value < 0.05. Identifications (IDs) of putative *Mtb*-derived peptides were rejected if any peptide-spectrum matches (PSMs) for the same peptide were found in the unfiltered DDA MS data for the corresponding mock-infected control. For each putative *Mtb* peptide identified, MS/MS spectra and extracted ion chromatograms (XIC) were manually inspected, and the ID was only accepted for further validation if it met the following criteria: (1) MS/MS spectra contained enough information to unambiguously assign a majority of the peptide sequence; (2) neutral losses were consistent with the chemical properties of the peptide; (3) manual *de novo* sequencing did not reveal an alternate peptide sequence that would explain a greater number of MS/MS spectrum peaks; (4) XIC showed a peak in MS intensity at the mass to charge ratio (m/z) of the peptide precursor ion at the retention time at which it was identified that did not appear in the corresponding mock-infected control. Peptides that met these criteria were further validated using SureQuant (see below).

DDA MS data were searched a second time against a database comprising only known T7SS substrates of *Mtb* H37Rv (all Esx-family proteins, all PE/PPE proteins, and EspA, EspB, EspC, EspE, EspF, EspI, EspJ, and EspK). Because high sequence similarity among many T7SS substrates artificially raises percolator q-values computed using this limited database, these searches were filtered only on ions score ≥ 20 rather than on ions score ≥ 15 and percolator q-value < 0.05. The only validated *Mtb* peptide identified via these additional searches that had not previously been identified (AEHGMPGVP, derived from PPE51) was omitted from enrichment analyses (see below) to avoid introducing bias.

### Gibbs clustering

Host peptides and putative *Mtb* peptides that passed the filters described above were clustered using GibbsCluster 2.0,^16^ hosted at (https://services.healthtech.dtu.dk/service.php?GibbsCluster-2.0). Default parameters for MHC-I peptide clustering were used, with the modification that a maximum of 6 clusters was allowed. The number of clusters that gave the highest KL divergence score was considered optimal. Clusters were assigned to HLA alleles by comparing cluster motifs to the known sequence binding motifs obtained from the NetMHCpan 4.1 motif viewer^53^ (https://services.healthtech.dtu.dk/service.php?NetMHCpan-4.1) for HLA alleles expressed by that donor.

### HLA allele assignment

For each donor, the binding affinity of *Mtb* peptides identified in DDA MS data for each HLA allele expressed by that donor (as determined by HLA typing − see above) was predicted using NetMHCpan 4.1.^53^ Peptides were assigned to the allele for which they had highest predicted binding score.

### Enrichment analyses

To determine whether *Mtb*-derived MHC-I peptides or their source proteins were enriched for proteins with specific secretion signals, we classified each protein in the reference *Mtb* H37Rv proteome (UP000001584) using SignalP 6.0,^54^ and added an additional class of known T7SS substrates (as defined above; see DDA data search). Under a “neutral model” in which all possible *Mtb* peptides are equally likely to be presented, the probability of drawing a peptide from a given class of proteins can be approximated as the sum of the lengths of the amino acid sequences of proteins in that class, divided by the total number of amino acid residues in the *Mtb* proteome. Using this approximation, we used a binomial test to assess whether the number of *Mtb*-derived peptides from each class of proteins detected in the MHC-I was greater or less than would be predicted under the neutral model. We used a hypergeometric test to assess whether the number of source proteins from each class was greater or less than expected, relative to a neutral model in which each protein in the *Mtb* proteome is equally likely to contribute to the MHC-I repertoire.

### SIL peptide synthesis

SIL peptides were synthesized at the MIT Biopolymers and Proteomics Lab using standard Fmoc chemistry using an Intavis model MultiPep peptide synthesizer with HATU activation and 5 μmol chemistry cycles. Starting resin used was Fmoc-Amide Resin (Applied Biosystems). Cleavage from resin and simultaneous amino acid side chain deprotection was accomplished using: trifluoroacetic acid (81.5% v/v); phenol (5% v/v); water (5% v/v); thioanisole (5% v/v); 1,2-ethanedithiol (2.5% v/v); 1% triisopropylsilane for 4 h. Standard Fmoc amino acids were procured from NovaBiochem and SIL Fmoc-Leu (^13^C_6_, ^15^N), Fmoc-Arg (^13^C_6_), Fmoc-Glu (^13^C_5_ ^15^N) were obtained from Cambridge Isotope Laboratories. Peptides were quality controlled by MSy and reverse phase chromatography using a Bruker MicroFlex MALDI-TOF and Agilent model 1100 HPLC system with a Vydac C18 column (300 Å, 5 μm, 2.1 × 150 mm^2^) at 300 μL/min monitoring at 210 and 280 nm with a trifluoroacetic acid/H_2_O/MeCN mobile phase survey gradient.

### Synthetic standard survey MS analyses

DDA MS analysis of the SIL peptide mixture was performed as described above (see DDA MS analyses) with the following modifications: Peptides were eluted using a gradient of 6-35% buffer B over 30 minutes, 35-45% over 2 minutes, 45-100% over 3 minutes, and 100% to 2% over 1 minute. No dynamic exclusion was used.

A second set of survey analyses was performed on the mixture of SIL peptides with background matrix using the full SureQuant acquisition method (see below). SIL peptides were spiked into a mixture of MHC-I peptides purified as described above from THP-1 cells differentiated into macrophages via 24 hours of treatment with 150 nM phorbol myristate acetate (PMA), which provided a representative background matrix. Because SIL amino acids are not 100% pure, SIL peptide concentrations were adjusted and survey analyses were repeated until the SIL peptide could be reliably detected while minimizing background signal detected at the mass of the biological peptide.

### SureQuant MS analyses

#### Validation analyses

Standard MS parameters and MS1 scan parameters were as described above (see DDA MS analyses). The custom SureQuant acquisition template available in Thermo Orbitrap Exploris Series 2.0 was used to build four methods for validation analyses. One method targeted peptides detected in DDA analyses of the MHC-I repertoire of macrophages from donors A, B, and C. The other three methods targeted peptides detected in DDA analyses of the MHC-I repertoire of macrophages from donors D, E, and F respectively. For each method, after the optimal charge state and most intense product ions were determined via a survey analysis of the synthetic SIL peptide standards (see above), one method branch was created for each m/z offset between the SIL peptide and biological peptide as previously described.^15^ A threshold of n = 3 out the top 6 product ions was used for pseudo-spectral matching, with a mass accuracy tolerance of 10 ppm.

#### Quantitative analyses

Quantitative SureQuant analyses were performed similarly, with an additional method branch added to target hipMHC peptides using an inclusion list. As none of the targets of these analyses has an m/z <380, an MS1 scan range of m/z 380-1500 was used to exclude some common background ions. Targeted MS data were analyzed using Skyline Daily Build 22.1.9.208. For each target peptide, the intensities of the three most intense product ions were integrated over the time during which the peptide was scanned. These intensities were normalized by the integrated intensities of the corresponding product ions from the corresponding SIL standard over the same time interval, and these ratios were averaged. Finally, these averaged ratios were normalized by the average ratio of the integrated intensities of the hipMHC standard peptides and normalized to the DMSO-treated, *Mtb*-infected condition to obtain the final relative abundance of each target. 100 fmol of SIL EsxA_28-36_ and 1 pmol of SIL EsxJKPW_24-34_ were spiked into each sample per analysis, as well as (where indicated) 250 fmol of SIL standard for each of the target self peptides. Self peptides were classified as derived from cytosolic/nuclear proteins or extracellular/endosomal proteins based on Uniprot annotations of the source protein(s). Peptides derived from transmembrane proteins were classified based on which side of the membrane they were predicted to reside on based on topological predictions made using DeepTMHMM.^55^

#### Immunofluorescence microscopy

100,000 macrophages per well plated on chamber slides (Ibidi) were infected with GFP-expressing *Mtb* at MOI 1 or mock-infected with media containing no *Mtb*. At the indicated time point post-infection, macrophages were washed with PBS and fixed with 4% paraformaldehyde (PFA) in PBS for 1 hour. After the full time course was collected, all slides were blocked with 5% normal goat serum in PBS supplemented with 0.3% v/v Triton X-100 for 1 hour at room temperature and stained overnight at 4°C with primary antibody [LAMP1 − Cell Signaling Technologies (CST) D2D11; P62 − CST D10E10; MHC-I − CST D8P1H; Galectin-3 − Thermo A3A12] diluted in antibody dilution buffer (PBS with 1% w/v bovine serum albumin and 0.3% v/v triton X-100). Wells were washed three times with PBS for 5 minutes each and stained with Alexa fluor 647 (AF647)-conjugated goat anti-rabbit or anti-mouse secondary antibody (Thermo) diluted to a final concentration of 1 μg/mL in antibody dilution buffer at room temperature for 2 hours. Wells were washed three times with PBS for 5 minutes each and stained with DAPI at a final concentration of 300 nM in PBS for 15 minutes at room temperature. Wells were washed three times with PBS for 5 minutes each, well dividers were removed, and slide covers were mounted on slides using Prolong Diamond anti-fade mounting media (Thermo). Slides were imaged using a TissueFAXS Confocal slide scanner system (TissueGnostics). 100 fields of view were acquired per condition using a 40x objective lens.

#### *Segmentation of* Mtb*-containing phagosomes*

Image segmentation was performed using opencv-python 4.5.4.60. A two-dimensional Gaussian blur with a standard deviation of 5 pixels was applied to de-noise GFP fluorescence images. A binary mask was generated from the blurred GFP image using a fluorescence intensity threshold that was empirically selected for each biological replicate. GFP+ phagosomes were further segmented using the watershed algorithm.

#### Colocalization analysis

A correlation image was generated by computing the Pearson correlation coefficient between the GFP fluorescent intensity and AF647 fluorescent intensity in a 41 × 41 pixel sliding window with a step size of 1. This correlation value was averaged over each *Mtb*-containing phagosome, and phagosomes with a mean value of greater than or equal to 0.6 were considered co-localized. For statistical analyses of log-transformed rates of co-localization, zeros were replaced with the limit of detection for a given sample [100% x (1/N) where N is the number of *Mtb*-containing phagosomes detected].

### *Mtb* growth curves

50,000 macrophages per well were infected in technical triplicate in opaque white 96-well plates with *luxABCDE-* expressing *Mtb* at MOI 2.5 (or mock-infected with media not containing *Mtb*). Where indicated, macrophages were treated with drug as described above, or media was supplemented with 10 ng/mL IFN-β (BioLegend) or 25 μg/mL rifampicin (RIF) after infection. Luminescence was measured at the indicated time points using a Spark 10M plate reader (Tecan).

### Phagocytosis assay

1 × 10^6^ macrophages were differentiated as described above on 6-well ultra low-attachment plates and infected with GFP-expressing *Mtb* at an MOI of 2.5 (or mock-infected with media containing no *Mtb*). After the 4-hour infection, macrophages were detached using PBS supplemented with 2 mM EDTA, washed with PBS, stained with Live/Dead Fixable Near-IR dye (Thermo) for 10 minutes, washed with FACS buffer (PBS supplemented with 2% FBS and 2 mM EDTA) and PBS, fixed with PBS containing 4% PFA for one hour, and washed and resuspended in FACS buffer for analysis on an LSRFortessa flow cytometer (BD).

### MHC-I flow cytometry

1 × 10^6^ macrophages were differentiated as described above on 6-well ultra low-attachment plates and infected with GFP-expressing *Mtb* at an MOI of 2.5 (or mock-infected with media containing no *Mtb*). 72 hours post-infection, macrophages were detached using PBS supplemented with 2 mM EDTA, blocked with Human TruStain FcX Fc receptor blocking solution (BioLegend) for 10 minutes, stained with phycoerythrin (PE)-conjugated anti-HLA-A,B,C antibody for 20 minutes (BioLegend, clone W6/32), stained with Live/Dead Fixable Near-IR dye (Thermo) for 10 minutes, washed with FACS buffer and PBS, fixed with PBS containing 4% PFA for one hour, and washed and resuspended in FACS buffer for analysis on an LSR Fortessa flow cytometer (BD).

### ELISA

50,000 macrophages per well were infected in technical triplicate in a 96-well plate format with *Mtb* at MOI 2.5 (or mock-infected with media not containing *Mtb*). 72 hours post-infection, culture media was collected and sterile filtered twice by centrifugation using 96-well 0.2 μm filter plates (Corning). The concentration of CXCL10 in the media was determined using a Human CXCL10 ELISA MAX kit (BioLegend 439904).

### LDH release assay

50,000 macrophages per well were infected in technical triplicate in a 96-well plate format with *Mtb* at MOI 2.5 (or mock-infected with media not containing *Mtb*) and treated with the indicated concentrations of drug or vehicle control. 24 hours post-infection, culture media was collected and sterile filtered twice by centrifugation using 96-well 0.2 μm filter plates (Corning). Percent cytotoxicity was determined using a CyQuant LDH Cytotoxicity Assay kit (Thermo C20300).

### Western blots

1 × 10^6^ macrophages were differentiated as described above on 6-well ultra low-attachment plates. Where indicated, macrophages were infected with *Mtb* at an MOI of 2.5. Macrophages were treated for 24 hours with the indicated concentration of MG-132 or DMSO control, and were then detached using PBS supplemented with 2 mM EDTA, washed with PBS, pelleted, and lysed in 100 μL RIPA buffer. Lysates were cleared by centrifugation at 16,000 x g for 5 minutes. For cells infected with *Mtb*, lysates were filtered twice using 96-well 0.2 μm filter plates (Corning).

NuPAGE LDS Sample Buffer (Thermo) and NuPAGE Sample Reducing Agent (Thermo) were added to a final concentration of 1x, and 20 μL of lysate per well was separated by SDS-PAGE. Protein was transferred to a PVDF membrane using an iBlot2 dry transfer system (Thermo). Membranes were blocked for 1 hour at room temperature with blocking buffer (Licor) and incubated overnight at 4°C with anti-K48-linked poly-ubiquitin antibody (Cell Signaling Technologies) and anti-β-actin antibody (Santa Cruz Biotechnologies), each diluted 1:1000 in antibody diluent (Licor). Blots were washed three times for 5 minutes each with tris-buffered saline with 0.1% tween-20 (TBS-T) and incubated for 1 hour at room temperature with IRDye 800CW goat anti-rabbit secondary antibody (Licor) and IRDye 680CW goat anti-mouse secondary antibody (Licor), each diluted 1:10,000 in antibody diluent (Licor). Blots were washed three times for five minutes each with TBS-T and imaged using an Odyssey DLx imaging system (Licor).

### Cathepsin B activity assays

Cathepsin B activity was measured using an assay protocol adapted from prior literature.^56,57^ 50,000 macrophages per well were infected in technical triplicate in a 96-well plate format with *Mtb* at MOI 2.5 (or mock-infected with media not containing *Mtb*) and treated with the indicated concentrations of E64d or vehicle control. 24 hours post-infection, macrophages were washed with PBS and lysed by adding 100 μL assay buffer (87.7 mM KH_2_PO_4_,12.3 mM NaHPO_4_, 4 mM EDTA, pH 5.5) supplemented with 2.6 mM dithiothreitol and 0.1% v/v Triton X-100, mixing thoroughly, and incubating at 37°C for 15 minutes. 40 μL of lysate was transferred to a separate 96-well plate, and the fluorogenic cathepsin B substrate Z-RR-AMC (Sigma-Aldrich) diluted in assay buffer was added to a final concentration of 2 mM. Plates were immediately transferred to a Spectramax iD3 plate reader (Molecular Devices) pre-warmed to 37°C.

Fluorescence at an absorbance wavelength of 370 nm and emission wavelength of 460 nm was measured in kinetic read mode at intervals of 2 minutes for a total duration of 20 minutes. The linear (enzyme-limited) regime of the fluorescence curve was determined to last approximately 10 minutes, so cathepsin B activity was quantified by using linear regression to determine the initial rate of fluorescence increase over the first 10 minutes of the assay.

### Lysotracker microscopy

100,000 macrophages per well plated on chamber slides (Ibidi) were infected with GFP-expressing *Mtb* at MOI 2.5 (or mock-infected with media containing no *Mtb*) and treated with bafilomycin or vehicle control as described above. 24 hours post-infection, macrophages were washed with Hank’s Balanced Salt Solution (HBSS), then incubated for 5 minutes at 37°C in pre-warmed HBSS containing 1 μM Lysotracker Red (Thermo). Cells were washed twice with HBSS and fixed with PBS containing 4% PFA for one hour, then stained with DAPI and mounted as described above (see immunofluorescence microscopy). Slides were imaged using a TissueFAXS Confocal slide scanner system (TissueGnostics). 100 fields of view were acquired per condition using a 40x objective lens.

#### Automated counting of nuclei

Image segmentation was performed using opencv-python 4.5.4.60. A two-dimensional Gaussian blur with a standard deviation of 5 pixels was applied to de-noise DAPI fluorescence images. A binary mask was generated from the smoothed DAPI image using a fluorescence intensity threshold that was determined automatically using Otsu’s thresholding method, as implemented in opencv-python. To disambiguate adjacent or overlapping nuclei, nuclei were further segmented using the watershed algorithm.

#### Calculation of Lysotracker+ area per cell

A two-dimensional Gaussian blur with a standard deviation of 5 pixels was applied to de-noise Lysotracker Red fluorescence images. A binary mask was generated from the Lysotracker Red image using a fixed intensity threshold for all samples. The total Lysotracker+ area was computed from the binary mask and divided by the number of nuclei counted in the image (see above).

### Curve fitting and statistical analyses

Unless otherwise indicated, all nonlinear curve fitting and all statistical tests were performed in GraphPad Prism 9.

## Acknowledgements

The authors thank all the staff members of the Ragon Institute, the Koch Institute, and MIT for the essential work they do to make our research possible. The authors thank Lauren Stopfer, Akeem Ngomu Akilimali, Charul Jani, Yong Xie (Ragon Institute BSL3 Facility), Thomas Diefenbach (Ragon Institute Microscopy Core Facility), Tigist Tamir, Cameron Flower, Angela Ahn, Kiera Clayton, and Aaron Shulkin for helpful conversations, training, and technical guidance. Iris Abrahantes Morales isolated monocytes from buffy coats. The authors thank Yuko Yuki and Mary Carrington (National Cancer Institute) for HLA genotyping of primary human cells. Kiera Clayton generously shared HLA-A*02:01+, HLA-B*57:01+ PBMCs from a leukapheresis sample for one biological replicate. The *eccCa1:Tn Mtb* strains used in this study were generously provided by Charul Jani and Amy Barczak. Diane Ballestas and Isadora Deese provided administrative support. Alla Leshinsky, Heather Amoroso, and Richard Cook (Koch Institute Biopolymers Core Facility) synthesized and purified SIL peptide standards. We adapted code written by Cameron Flower to plot MS/MS spectra. The authors thank the staff of the Ragon Institute Flow Cytometry Core Facility. This work was supported by funding from NIH grants R35GM142900-01 and R01A1022553, NIEHS grant P42 ES027707, and the Center for Precision Cancer Medicine.

## Author Contributions

O.L. performed experiments and analyzed data. B.D.B., F.M.W., and O.L. conceptualized and designed experiments. B.D.B. and F.M.W. acquired funding and supervised the work. O.L. wrote the initial manuscript draft. B.D.B., F.M.W., and O.L. revised the manuscript.

