## Supplemental Figures and Tables for "Immunopeptidomics reveals determinants of *Mycobacterium tuberculosis* antigen presentation on MHC class I"

**Supplementary Table 1. Class I HLA alleles expressed by primary monocyte donors.**

| Donor | HLA-A allele 1 | HLA-A allele 2 | HLA-B allele 1 | HLA-B allele 2 | HLA-C allele 1 | HLA-C allele 2 |
|---|---|---|---|---|---|---|
| A | ND | ND | ND | ND | ND | ND |
| B | 02:01 | 24:02 | 07:02 | 27:07 | 07:02 | 15:02 |
| C | 01:01 | 02:01 | 44:03 | 57:01 | 06:02 | 16:01 |
| D | 23:01 | 26:01 | 35:01 | 50:01 | 04:01 | 06:02 |
| E | 11:01 | 24:02 | 39:06 | 51:01 | 07:02 | 15:02 |
| F | 01:01 | 26:01 | 35:01 | 38:01 | 04:01 | 12:03 |

**Supplementary Table 2. *Mtb*-derived peptides that passed manual inspection of DDA MS data but failed SureQuant validation.**

| Source protein | Peptide sequence |
|---|---|
| Rv0383c | AAPGRPVAPG |
| pyrD | GDRLALISV |
| Rv2303c | KHPNVYLEL |
| Rv0839 | YTHGYHES |
| kgd | AERAAAAAP |
| Rv1375 | EAAQSRITA |
| GabD1 | AKVGASAAY |
| PE1 | AAGNLRAAI |
| HlfX | IPYDRGDLV |
| Rv2807 | AKWILEGIK |
| Rv3818 | IAPELVRT |
| Rv1065 | YTRIHGDEEL |
| Rv0293c | DELIAGLAY |
| Rv3779 | VAIAVGPALT |
| PPE55 | TVAPINLNP |
| Rv2263 | QEIEEGIL |
| Rv0333 | GEDPGIAR |

**Supplementary Figure 1. Performance of two BSL-3 compatible MHC-I immunopeptidomics workflows.** a) Length distribution of MHC-I peptides identified by MS for protocol 1 and protocol 2 (see Figure 1a and Methods), for all detected peptides and validated *Mtb*-derived peptides. b) Predicted hydrophobicity index (computed using the ProtViz R package; https://rdrr.io/cran/protViz/man/ssrc.html) as a function of retention time for protocols 1 and 2. Green points indicate validated *Mtb* peptides. c) Absolute *Mtb* protein abundance in axenic culture as a function of protein rank, as quantified by Schubert et al.[27] Red points indicate source proteins of peptides detected in the MHC-I repertoire of *Mtb*-infected macrophages (see Figure 1c).

**Supplementary Figure 2. Gibbs clustering groups MHC-I peptides into clusters that correspond to HLA alleles expressed by each donor.** Heatmap of peptide binding score predicted by NetMHCpan 4.1, Gibbs clustering motif, and corresponding published binding motif for each HLA allele expressed by a) donor B, b) donor C, c) donor D, d) donor E, e) donor F.

**Supplementary Figure 3. Predicted class I HLA binding of *Mtb*-derived MHC-I peptides.** Heatmap of peptide binding score predicted by NetMHCpan 4.1 for *Mtb*-derived peptides for each HLA allele expressed by a) donor B, b) donor C, c) donor D, d) donor E, e) donor F. Label colors correspond to the clusters defined by Gibbs clustering of the full set of identified peptides (see Supplementary Figure 2).

**Supplementary Figure 4. Workflow for validating *Mtb*-derived MHC-I peptide identifications by SureQuant.** Schematic showing how MHC-I peptide samples were divided between DDA-based discovery analyses (75%) and SureQuant-based validation analyses (25%). MS/MS spectra from survey analyses of mixtures of SIL peptide standards alone were compared to the MS/MS spectra of the corresponding biological peptides. Chromatograms demonstrating co-elution of biological peptides with the corresponding SIL standards were obtained by SureQuant.

**Supplementary Figure 5. Validation of *Mtb*-derived MHC-I peptide identifications.** Representative MS/MS spectrum comparisons (left column) and SureQuant product ion chromatograms for *Mtb*-infected and mock-infected macrophages (right column) for each peptide. For each peptide, the amino acid that is stable isotope labeled in the synthetic standard is indicated in red.

**Supplementary Figure 6. Immunofluorescence staining for Galectin-3, P62, LAMP-1, and MHC-I is specific**. Representative spinning disk confocal microscopy images comparing macrophages stained with primary antibody specific for galectin-3 (a), P62 (b), LAMP-1 (c), or MHC-I (d) to those stained with secondary antibody alone, 120h post-infection.

**Supplementary Figure 7. Infection of macrophages with wild-type or ESX-1-deficient *Mtb* strains results in similar rates of infection and does not affect surface MHC-I levels 72 hours post-infection.** Macrophages infected with GFP-expressing wild-type (H37Rv) or ESX-1-deficient (*eccCa1:Tn*) *Mtb* were surface stained for MHC-I 72 hours post-infection and analyzed by flow cytometry. a) Representative histograms of GFP fluorescence intensity vs. autofluorescence, showing gating for GFP+ infected cells. b) Proportion of GFP+ cells for n=3 donors (** $p < 0.01$, *** $p < 0.001$; one-way ANOVA with Tukey's multiple comparisons test). c) Representative histograms of surface MHC-I fluorescence intensity. d) Mean MHC-I fluorescence intensity (MFI) for n=3 donors (p-values determined by one-way ANOVA with Tukey's multiple comparisons test).

**Supplementary Figure 8. MG-132, E64d, and bafilomycin effectively inhibit their targets in *Mtb*-infected macrophages.** Where applicable, red arrows indicate the dose of each drug used in MS experiments. a) Representative western blots for K48-linked polyubiquitinated protein in uninfected or *Mtb*-infected macrophages. β-actin is included as a loading control. b) Quantification of K48-polyubiquitinated protein by densitometry of western blots for n=3 donors. Values are normalized to the abundance of β-actin, also quantified by densitometry of western blots. EC50 was determined by fitting a curve of the form 100% × [MG-132]/([MG-132] + EC50). Data points and error bars represent the mean ± standard deviation. c) Relative cathepsin B activity (as measured via hydrolysis of the fluorogenic cathepsin B substrate Z-RR-AMC; see Methods) as a function of E64d concentration in mock-infected and *Mtb*-infected macrophages. EC50 was determined by fitting a curve of the form 100% × [1 - [E64d]/([E64d] + EC50)]. Data points and error bars represent the mean ± standard deviation. d) Representative spinning disk confocal microscopy images of mock-infected or *Mtb*-infected macrophages treated with 10 nM bafilomycin for 24 hours and stained with lysotracker red. e) Mean lysotracker-positive area per cell for n = 3 donors. (* $p < 0.05$, *** $p < 0.001$; paired one-tailed T-test).

**Supplementary Figure 9. MG-132, E64d, and bafilomycin have minimal cytotoxicity at effective doses in *Mtb*-infected macrophages.** Mock-infected or *Mtb*-infected macrophages were treated with each drug at the indicated concentrations, and the change in cell viability was measured by lactate dehydrogenase (LDH) release assay. Where applicable, red arrows indicate the dose of each drug used in MS experiments. Data points and error bars represent the mean ± standard deviation. a) MG-132 (n = 4 donors), b) bafilomycin (10 nM; n = 4 donors), c) E64d (n = 3 donors).

**Supplementary Figure 10. Treatment of macrophages with MG-132, E64d, and bafilomycin does not impair phagocytosis or outgrowth of *Mtb*.** Macrophages were pre-treated with media containing drug or vehicle control for 1 hour and infected with *Mtb* in media containing drug or vehicle control, as in MS experiments. a) Representative flow cytometry plots of GFP fluorescence vs. autofluorescence in macrophages infected with GFP-expressing *Mtb*, immediately after being allowed to phagocytose *Mtb* for 4 hours at an MOI of 2.5. b) Quantification of the proportion of macrophages that took up *Mtb* (i.e., were GFP+) (p-values determined by one-way ANOVA with Dunnett's multiple comparisons test). c) Luminescence as a function of time post-infection for macrophages infected with *Mtb* expressing a luciferase reporter. Rifampicin (RIF) is included as a control demonstrating inhibition of growth. Datapoints represent the mean ± standard deviation for n = 3 donors, each representing the mean of three technical replicates.

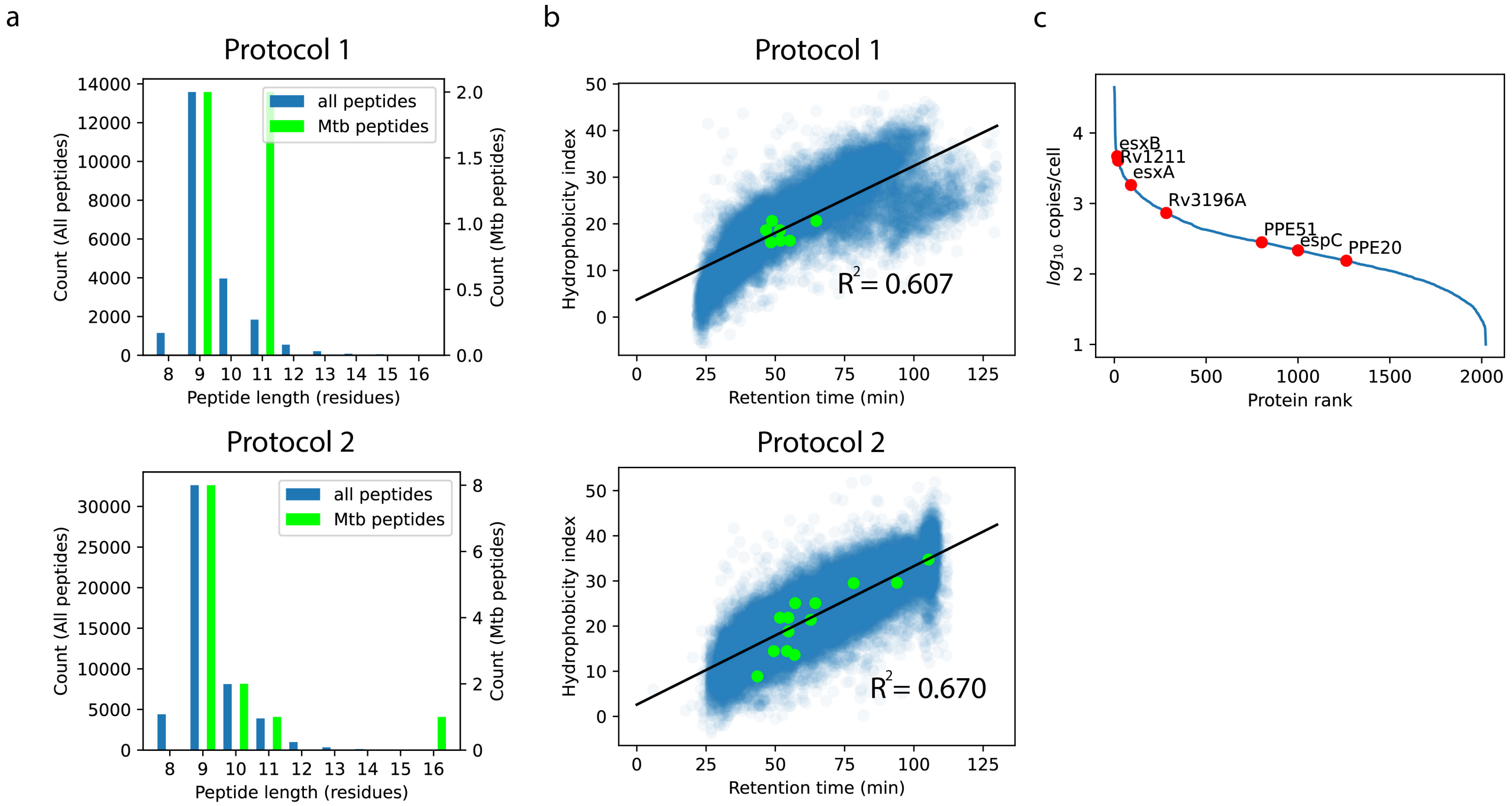

a
Protocol 1
all peptides
Mtb peptides
Count (All peptides)
Count (Mtb peptides)
Peptide length (residues)
Protocol 2
b
Protocol 1
Hydrophobicity index
Retention time (min)
$R^2$ = 0.607
Protocol 2
$R^2$ = 0.670
c
$log_{10}$ copies/cell
Protein rank
esxB
Rv1211
esxA
Rv3196A
PPE51
espC
PPE20

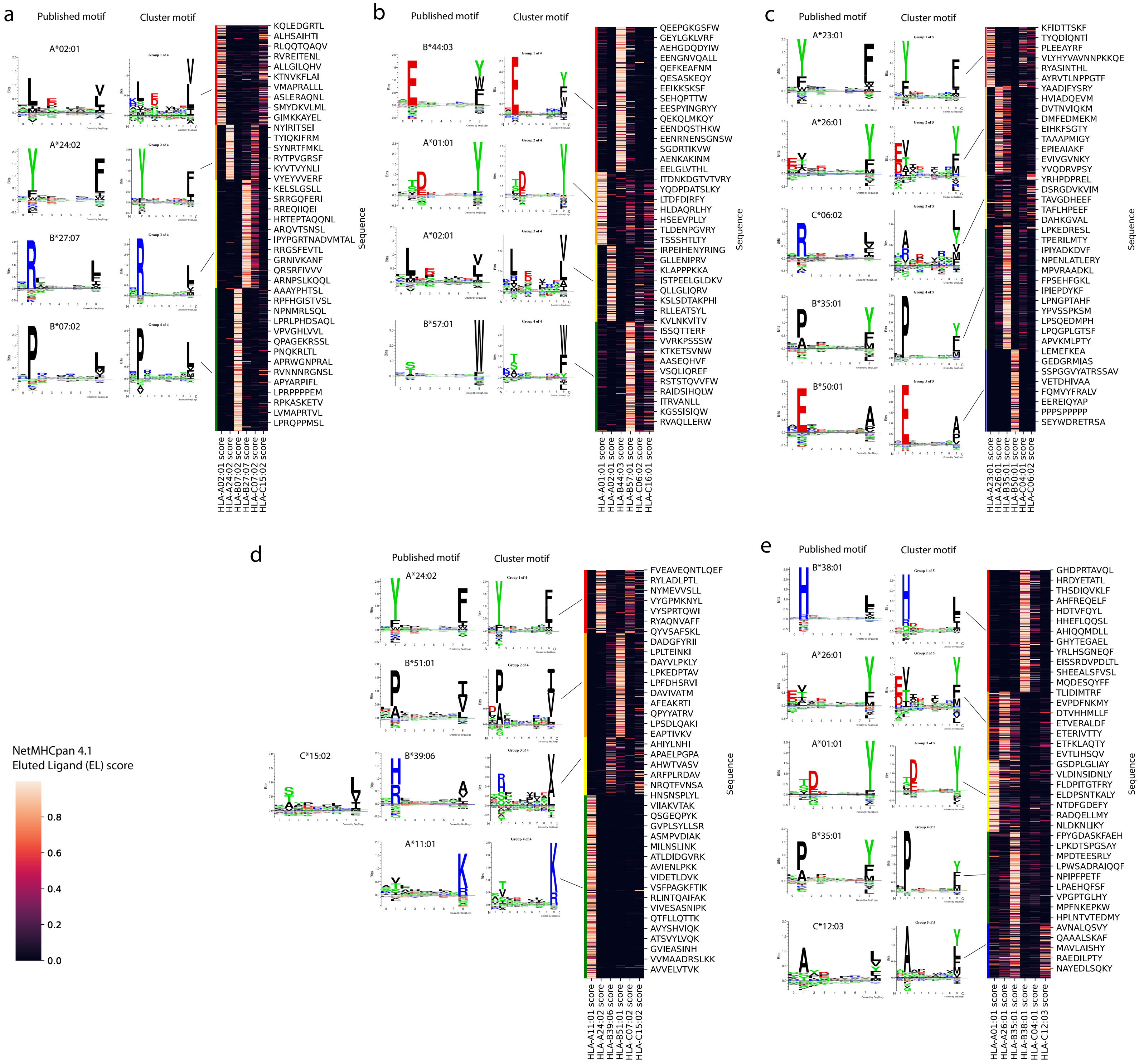
a
Published motif
Cluster motif
A*02:01
A*24:02
B*27:07
B*07:02
Group 1 of 4
Group 2 of 4
Group 3 of 4
Group 4 of 4
KQLEDGRTL
ALHSAIHTI
RLQQTQAQV
RVREITENL
ALLGILQHV
KTNVKFLAI
VMAPRALLL
ASLERAQNL
SMYDKVLML
GIMKKAYEL
NYIRITSEI
TYIQKIFRM
SYNRTFMKL
RYTPVGRSF
KYVTVYNLI
VYEYVVERF
KELSLGSLL
SRRGQFERI
RREQIIQEI
HRTEPTAQQNL
ARQVTSNSL
IPYPGRTNADVMTAL
RRGSFEVTL
GRNIVKANF
QRSRFIVVV
ARNPSLKQQL
AAAYPHTSL
RPFHGISTVSL
NPNMRLSQL
LPRLPHDSAQL
VPVGHLVVL
QPAGEKRSSL
PNQKRLTL
APRWGNPRAL
RVNNNRGNSL
APYARPIFL
LPRPPPPEM
RPKASKETV
LVMAPRTVL
LPRQPPMSL
Sequence
HLA-A02:01 score
HLA-A24:02 score
HLA-B07:02 score
HLA-B27:07 score
HLA-C07:02 score
HLA-C15:02 score
b
Published motif
Cluster motif
B*44:03
A*01:01
A*02:01
B*57:01
QEEPGKGSFW
GEYLGKLVRF
AEHGDQDYIW
EENGNVQALL
QEFKEAFNM
QESASKEQY
EEIKKSKSF
SEHQPTTW
EESPYINGRYY
QEKQLMKQY
EENDQSTHKW
EENRNENSGNSW
SGDRTIKVW
AENKAKINM
EELGLVTHL
ITDNKDGTVTVRY
YQDPDATSLKY
LTDFDIRFY
HLDAQRLHY
HSEEVPLLY
TLDENPGVRY
TSSSHTLTY
IRPEIHENYRING
GLLENIPRV
KLAPPPKKA
ISTPEELGLDKV
QLLGLIQRV
KSLSDTAKPHI
RLLEATSYL
KVLNKVITV
ISSQTTERF
VVRKPSSSW
KTKETSVNW
AASEQHVF
VSQLIQREF
RSTSTQVVFW
RAIDSIHQLW
ITRVANLL
KGSSISIQW
RVAQLLERW
Sequence
HLA-A01:01 score
HLA-A02:01 score
HLA-B44:03 score
HLA-B57:01 score
HLA-C06:02 score
HLA-C16:01 score
c
Published motif
Cluster motif
A*23:01
A*26:01
C*06:02
B*35:01
B*50:01
KFIDTTSKF
TYQDIQNTI
PLEEAYRF
VLYHYVAVNNPKKQE
RYASINTHL
AYRVTLNPPGTF
YAADIFYSRY
HVIADQEVM
DVTNVIQKM
DMFEDMEKM
EIHKFSGTY
TAAAPMIGY
EPIEAIAKF
EVIVGVNKY
YVQDRVPSY
YRHPDPREL
DSRGDVKVIM
TAVGDHEEF
TAFLHPEEF
DAHKGVAL
LPKEDRESL
TPERILMTY
IPIYADKDVF
NPENLATLERY
MPVRAADKL
FPSEHFGKL
IPIEPDYKF
LPNGPTAHF
YPVSSPKSM
LPSQEDMPH
LPQGPLGTSF
APVKMLPTY
LEMEFKEA
GEDGRMIAS
SSPGGVYATRSSAV
VETDHIVAA
FQMVYFRALV
EEREIQYAP
PPPSPPPPP
SEYWDRETRSA
Sequence
HLA-A23:01 score
HLA-A26:01 score
HLA-B35:01 score
HLA-B50:01 score
HLA-C04:01 score
HLA-C06:02 score
d
Published motif
Cluster motif
C*15:02
A*24:02
B*51:01
B*39:06
A*11:01
FVEAVEQNTLQEF
RYLADLPTL
NYMEVVSLL
VYGPMKNYL
VYSPRTQWI
RYAQNVAFF
QYVSAFSKL
DADGFYRII
LPLTEINKI
DAYVLPKLY
LPKEDPTAV
LPFDHSRVI
DAVIVATM
AFEAKRTI
QPYYATRV
LPSDLQAKI
EAPTIVKV
AHIYLNHI
APAELPGPA
AHWTVASV
ARFPLRDAV
NRQTFVNSA
HNSNSPLYL
VIIAKVTAK
QSGEQPYK
GVPLSYLLSR
ASMPVDIAK
MILNSLINK
ATLDIDGVRK
AVIENLPKK
VIDETLDVK
VSFPAGKFTIK
RLINTQAIFAK
VIVESASNIPK
QTFLLQTTK
AVYSHVIQK
ATSVYLVQK
GVIEASINH
VVMAADRSLKK
AVVELVTVK
Sequence
HLA-A11:01 score
HLA-A24:02 score
HLA-B39:06 score
HLA-B51:01 score
HLA-C07:02 score
HLA-C15:02 score
NetMHCpan 4.1
Eluted Ligand (EL) score
0.8
0.6
0.4
0.2
0.0
e
Published motif
Cluster motif
B*38:01
A*26:01
A*01:01
B*35:01
C*12:03
GHDPRTAVQL
HRDYETATL
THSDIQVKLF
AHFREQELF
HDTVFQYL
HHEFLQQSL
AHIQQMDLL
GHYTEGAEL
YRLHSGNEQF
EISSRDVPDLTL
SHEEALSFVSL
MQDESQYFF
TLIDIMTRF
EVPDFNKMY
DTVHHMLLF
ETVERALDF
ETERIVTTY
ETFKLAQTY
EVTLIHSQV
GSDPLGLIAY
VLDINSIDNLY
FLDPITGTFRY
ELDPSNTKALY
NTDFGDEFY
RADQELLMY
NLDKNLIKY
FPYGDASKFAEH
LPKDTSPGSAY
MPDTEESRLY
LPWSADRAIQQF
NPIPFPETF
LPAEHQFSF
VPGPTGLHY
MPFNKEPKW
HPLNTVTEDMY
AVNALQSVY
QAAALSKAF
MAVLAISHY
RAEDILPTY
NAYEDLSQKY
Sequence
HLA-A01:01 score
HLA-A26:01 score
HLA-B35:01 score
HLA-B38:01 score
HLA-C04:01 score
HLA-C12:03 score

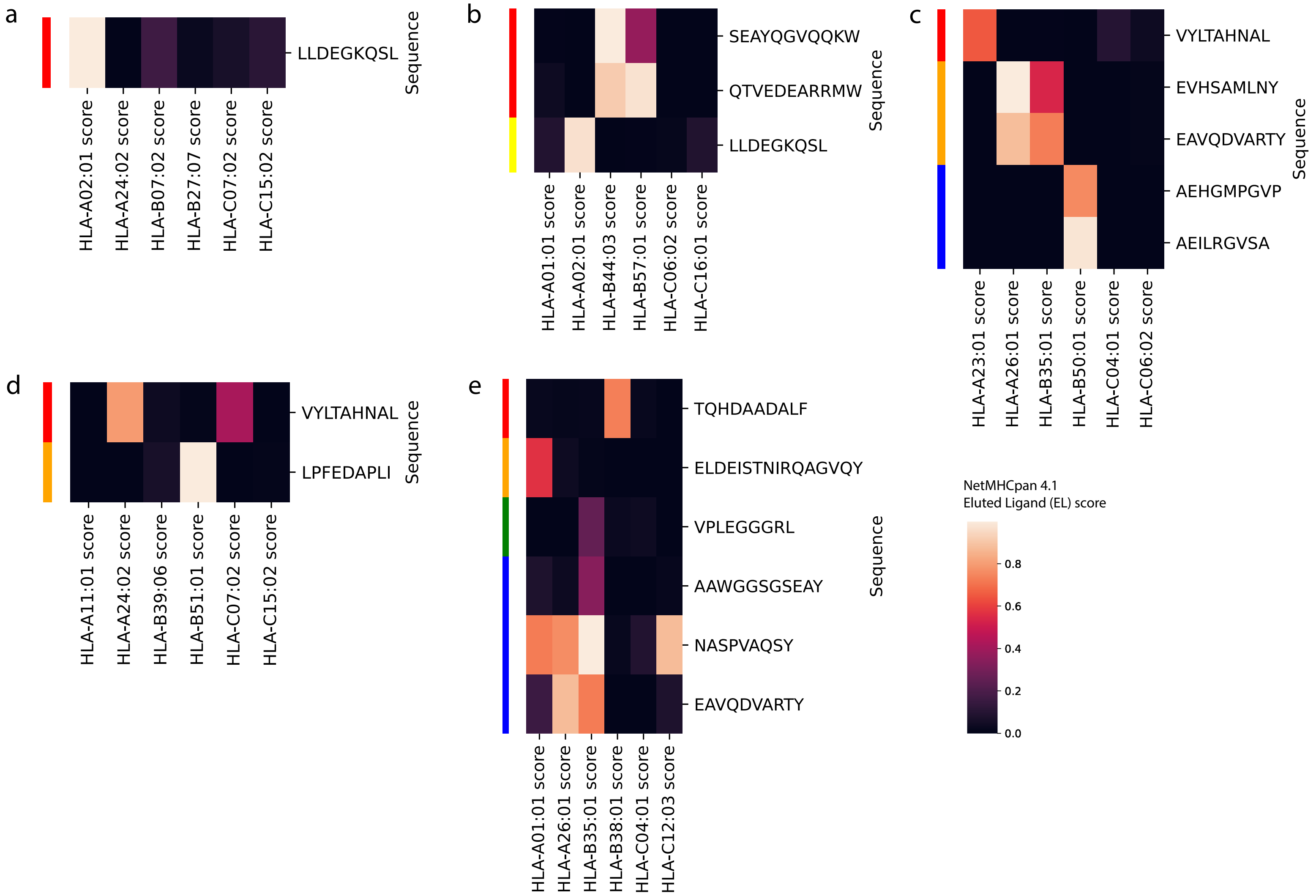
a
LLDEGKQSL
Sequence
HLA-A02:01 score
HLA-A24:02 score
HLA-B07:02 score
HLA-B27:07 score
HLA-C07:02 score
HLA-C15:02 score
b
SEAYQGVQQKW
QTVEDEARRMW
LLDEGKQSL
Sequence
HLA-A01:01 score
HLA-A02:01 score
HLA-B44:03 score
HLA-B57:01 score
HLA-C06:02 score
HLA-C16:01 score
c
VYLTAHNAL
EVHSAMLNY
EAVQDVARTY
AEHGMPGVP
AEILRGVSA
Sequence
HLA-A23:01 score
HLA-A26:01 score
HLA-B35:01 score
HLA-B50:01 score
HLA-C04:01 score
HLA-C06:02 score
d
VYLTAHNAL
LPFEDAPLI
Sequence
HLA-A11:01 score
HLA-A24:02 score
HLA-B39:06 score
HLA-B51:01 score
HLA-C07:02 score
HLA-C15:02 score
e
TQHDAADALF
ELDEISTNIRQAGVQY
VPLEGGGRL
AAWGGSGSEAY
NASPVAQSY
EAVQDVARTY
Sequence
HLA-A01:01 score
HLA-A26:01 score
HLA-B35:01 score
HLA-B38:01 score
HLA-C04:01 score
HLA-C12:03 score
NetMHCpan 4.1
Eluted Ligand (EL) score
0.8
0.6
0.4
0.2
0.0

a

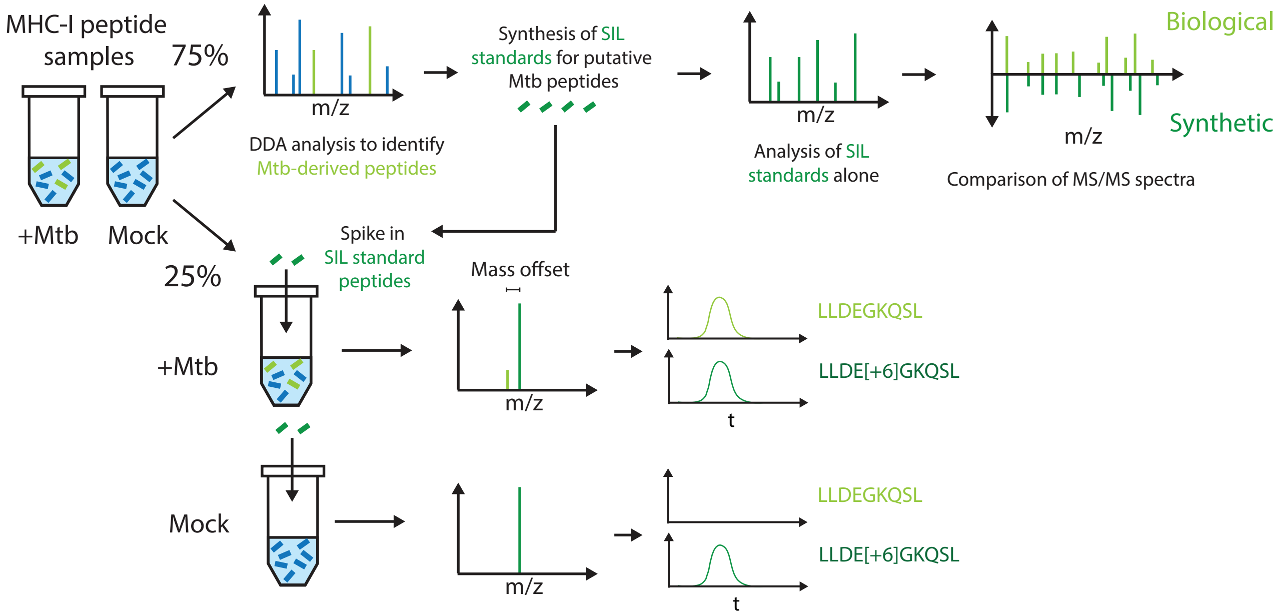
MHC-I peptide samples
75%
25%
+Mtb
Mock
m/z
DDA analysis to identify Mtb-derived peptides
Synthesis of SIL standards for putative Mtb peptides
m/z
Analysis of SIL standards alone
Biological
Synthetic
m/z
Comparison of MS/MS spectra
Spike in SIL standard peptides
Mass offset
+Mtb
m/z
LLDEGKQSL
LLDE[+6]GKQSL
t
Mock
m/z
LLDEGKQSL
LLDE[+6]GKQSL
t

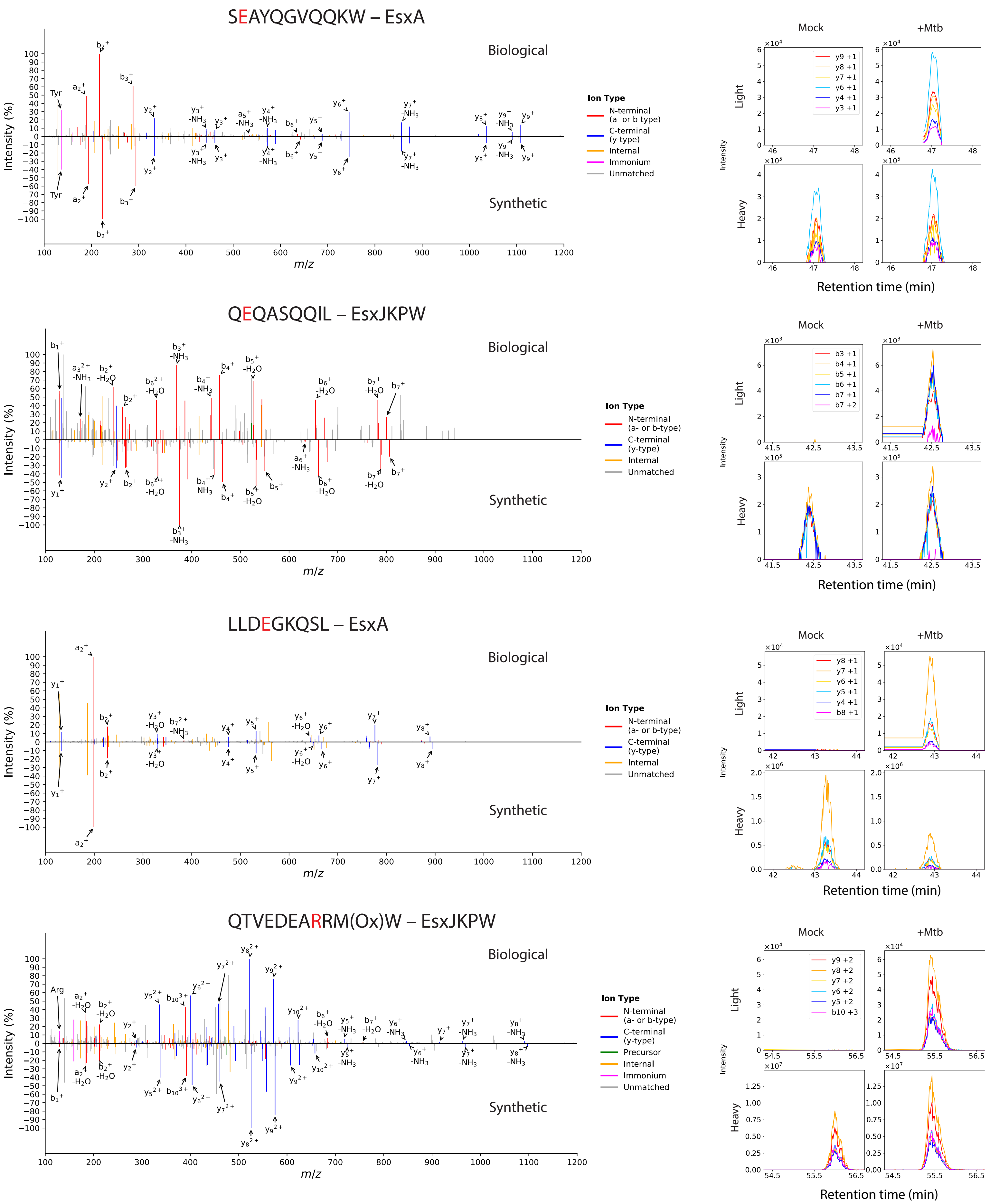
SEAYQGVQQKW – EsxA
QEQASQQIL – EsxJKPW
LLDEGKQSL – EsxA
QTVEDEARRM(Ox)W – EsxJKPW
Biological
Synthetic
Intensity (%)
m/z
Ion Type
N-terminal (a- or b-type)
C-terminal (y-type)
Precursor
Internal
Immonium
Unmatched
Mock
+Mtb
Light
Heavy
Intensity
Retention time (min)

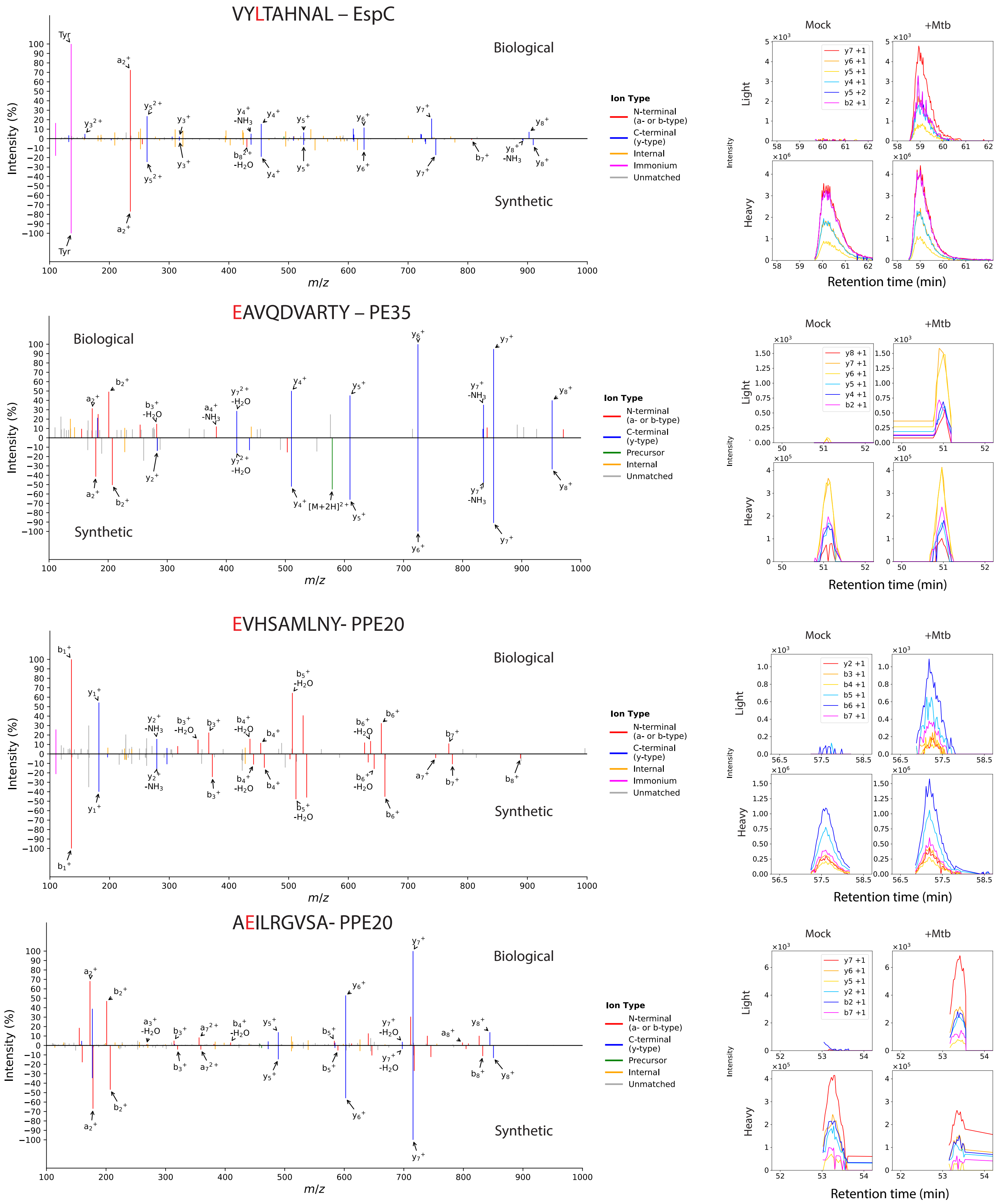
VYLTAHNAL – EspC
Biological
Synthetic
Intensity (%)
m/z
Ion Type
N-terminal (a- or b-type)
C-terminal (y-type)
Internal
Immonium
Unmatched
Mock
+Mtb
Light
Heavy
Intensity
Retention time (min)
EAVQDVARTY – PE35
Biological
Synthetic
Ion Type
N-terminal (a- or b-type)
C-terminal (y-type)
Precursor
Internal
Unmatched
Mock
+Mtb
Light
Heavy
Retention time (min)
EVHSAMLNY- PPE20
Biological
Synthetic
Ion Type
N-terminal (a- or b-type)
C-terminal (y-type)
Internal
Immonium
Unmatched
Mock
+Mtb
Light
Heavy
Retention time (min)
AEILRGVSA- PPE20
Biological
Synthetic
Ion Type
N-terminal (a- or b-type)
C-terminal (y-type)
Precursor
Internal
Unmatched
Mock
+Mtb
Light
Heavy
Retention time (min)

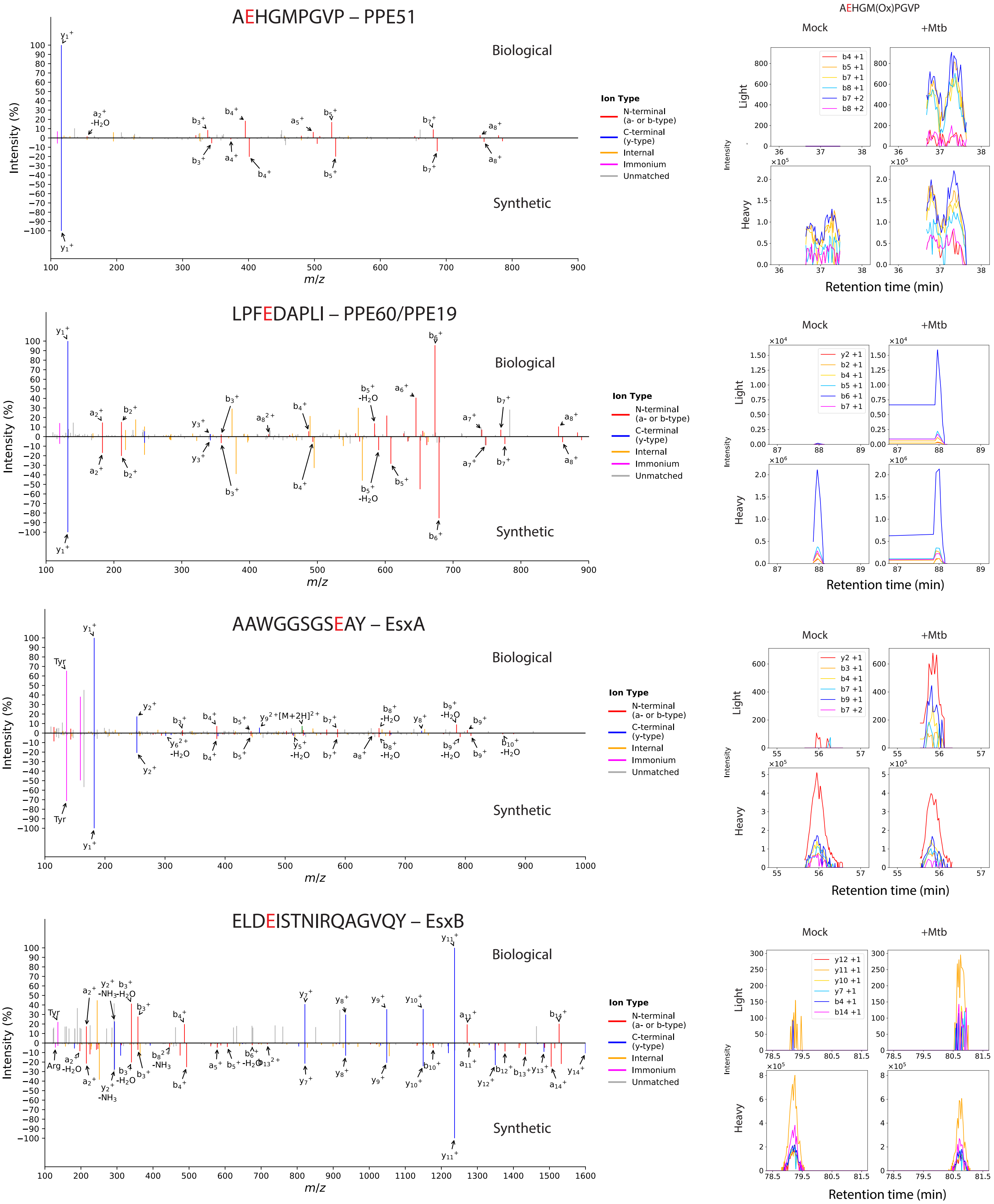

AEHGMPGVP – PPE51
Biological
Synthetic
Intensity (%)
m/z
Ion Type
N-terminal (a- or b-type)
C-terminal (y-type)
Internal
Immonium
Unmatched
AEHGM(Ox)PGVP
Mock
+Mtb
Light
Heavy
Intensity
b4 +1
b5 +1
b7 +1
b8 +1
b7 +2
b8 +2
Retention time (min)
LPFEDAPLI – PPE60/PPE19
Biological
Synthetic
Intensity (%)
m/z
Ion Type
N-terminal (a- or b-type)
C-terminal (y-type)
Internal
Immonium
Unmatched
Mock
+Mtb
Light
Heavy
Intensity
y2 +1
b2 +1
b4 +1
b5 +1
b6 +1
b7 +1
Retention time (min)
AAWGGSGSEAY – EsxA
Biological
Synthetic
Intensity (%)
m/z
Ion Type
N-terminal (a- or b-type)
C-terminal (y-type)
Internal
Immonium
Unmatched
Mock
+Mtb
Light
Heavy
Intensity
y2 +1
b3 +1
b4 +1
b7 +1
b9 +1
b7 +2
Retention time (min)
ELDEISTNIRQAGVQY – EsxB
Biological
Synthetic
Intensity (%)
m/z
Ion Type
N-terminal (a- or b-type)
C-terminal (y-type)
Internal
Immonium
Unmatched
Mock
+Mtb
Light
Heavy
Intensity
y12 +1
y11 +1
y10 +1
y7 +1
b4 +1
b14 +1
Retention time (min)

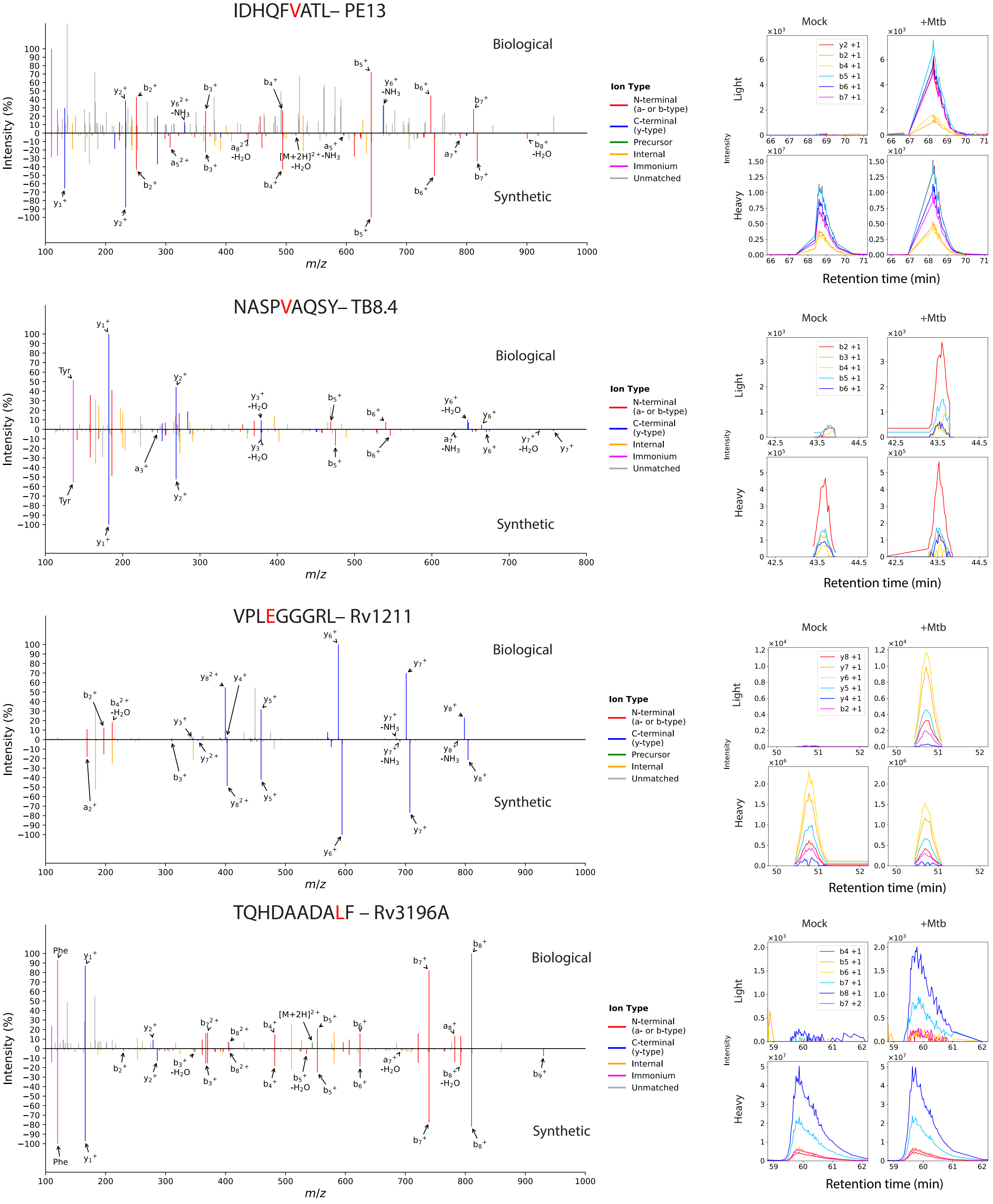

IDHQFVATL– PE13
NASPVAQSY– TB8.4
VPLEGGGRL– Rv1211
TQHDAADALF – Rv3196A
Biological
Synthetic
Intensity (%)
m/z
Ion Type
N-terminal (a- or b-type)
C-terminal (y-type)
Precursor
Internal
Immonium
Unmatched
Mock
+Mtb
Light
Heavy
Intensity
Retention time (min)
y2 +1
b2 +1
b4 +1
b5 +1
b6 +1
b7 +1
b3 +1
y8 +1
y7 +1
y6 +1
y5 +1
y4 +1
b8 +1
b7 +2
Tyr
Phe
[M+2H]$^{2+}$
[M+2H]$^{2+}$-NH$_3$-H$_2$O

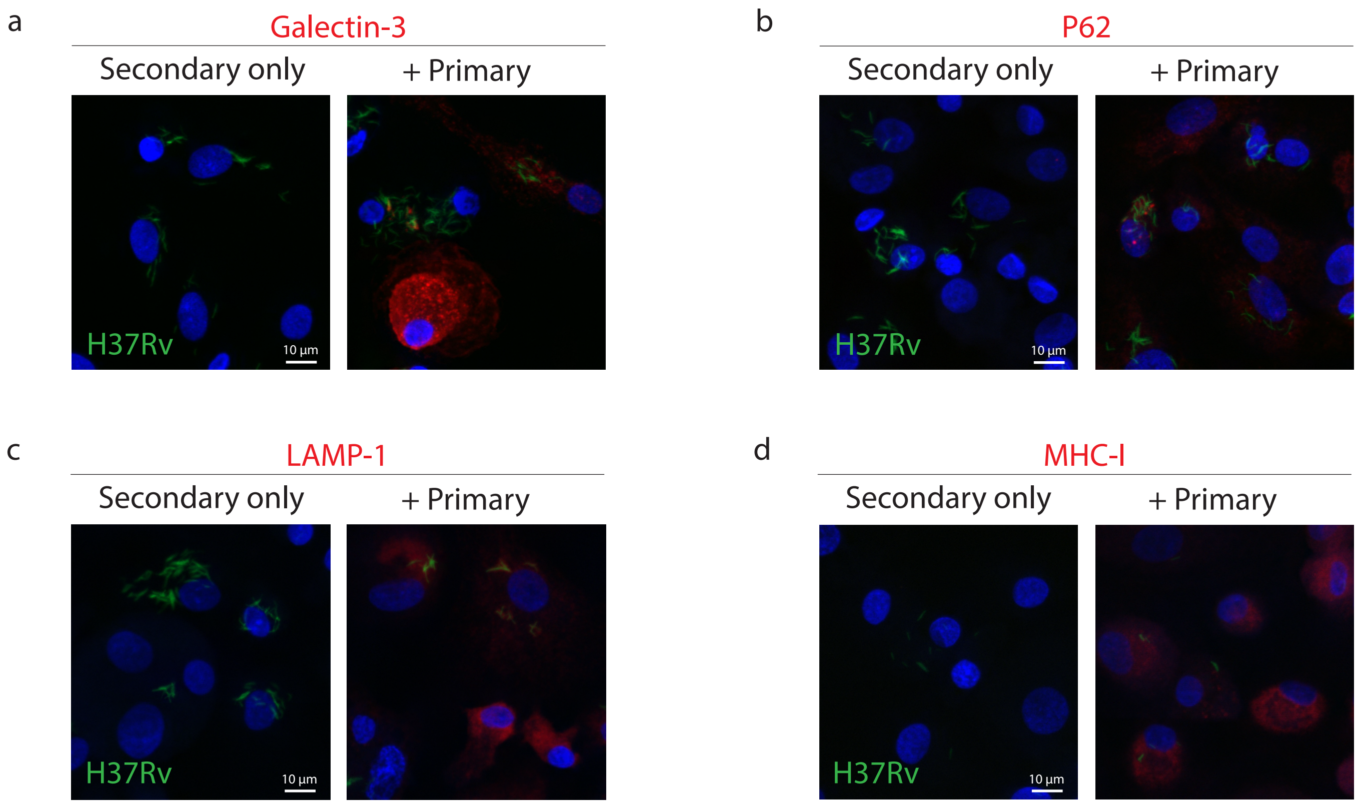
a
Galectin-3
Secondary only
+ Primary
H37Rv
10 µm
b
P62
Secondary only
+ Primary
H37Rv
10 µm
c
LAMP-1
Secondary only
+ Primary
H37Rv
10 µm
d
MHC-I
Secondary only
+ Primary
H37Rv
10 µm

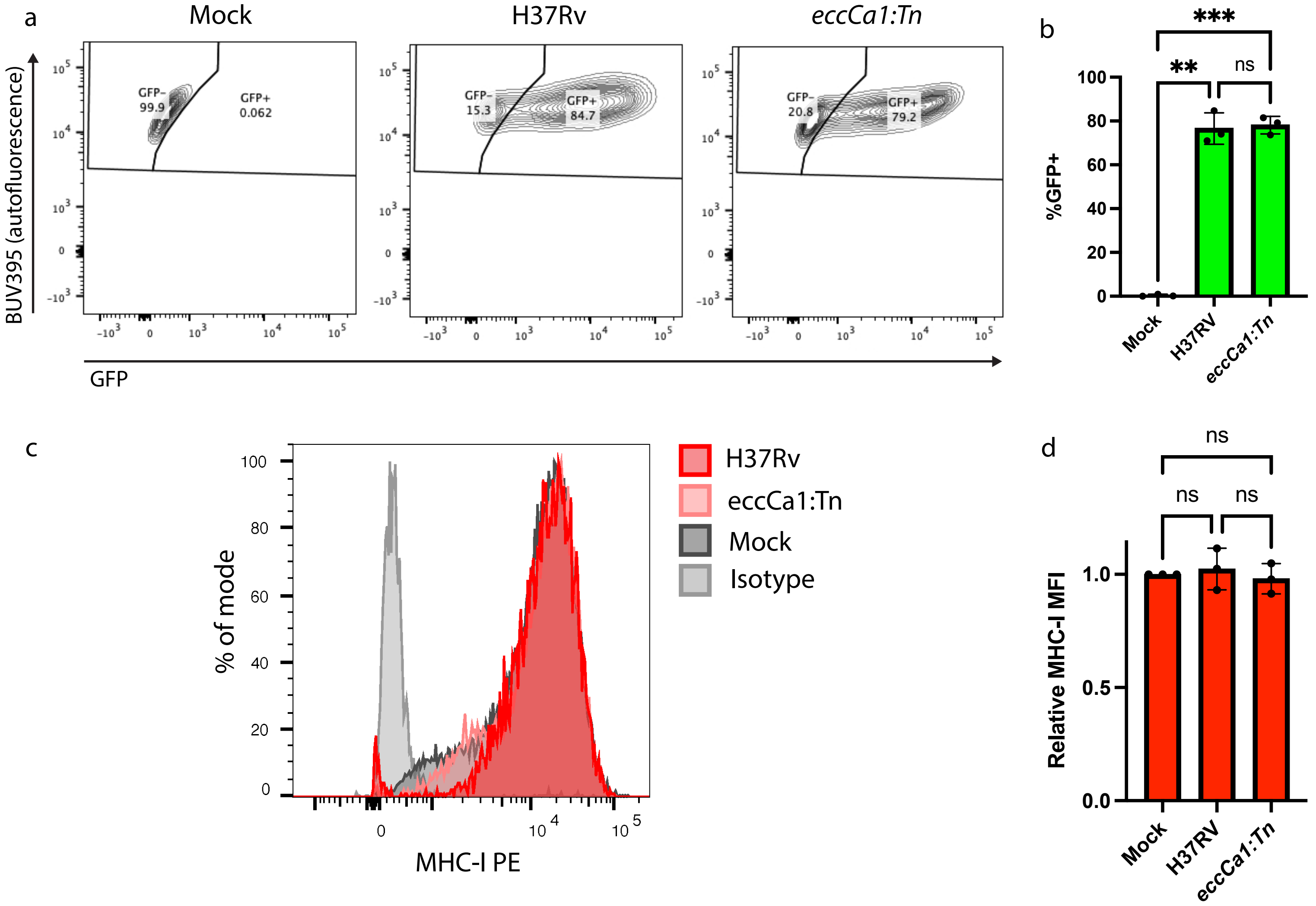
a
Mock
H37Rv
eccCa1:Tn
BUV395 (autofluorescence)
GFP
GFP-
99.9
GFP+
0.062
GFP-
15.3
GFP+
84.7
GFP-
20.8
GFP+
79.2
b
%GFP+
***
**
ns
Mock
H37RV
eccCa1:Tn
c
H37Rv
eccCa1:Tn
Mock
Isotype
% of mode
MHC-I PE
d
ns
ns
ns
Relative MHC-I MFI
Mock
H37RV
eccCa1:Tn

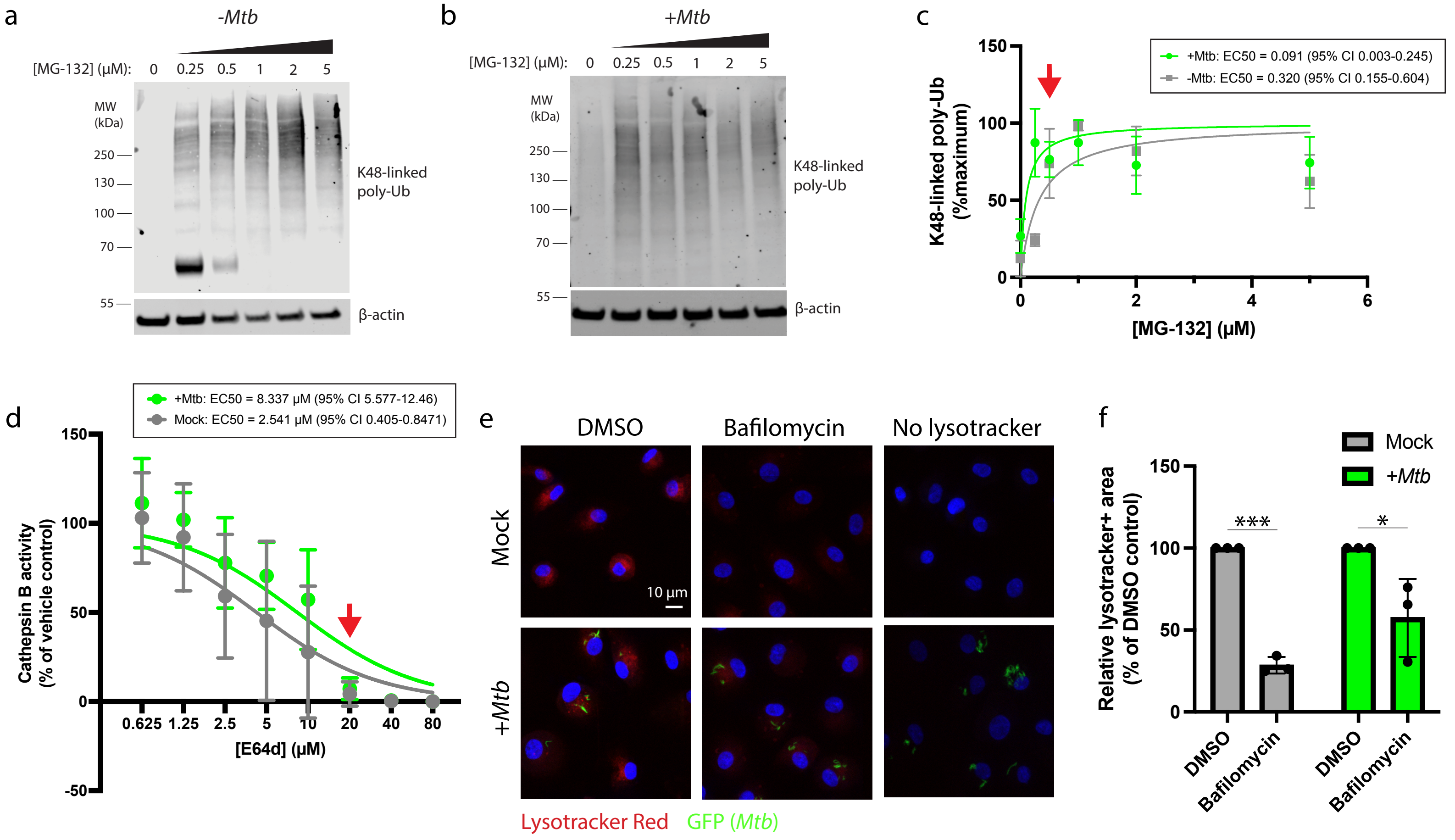

a
-Mtb
[MG-132] (μM): 0 0.25 0.5 1 2 5
MW (kDa)
250
130
100
70
55
K48-linked poly-Ub
β-actin
b
+Mtb
[MG-132] (μM): 0 0.25 0.5 1 2 5
MW (kDa)
250
130
100
70
55
K48-linked poly-Ub
β-actin
c
+Mtb: EC50 = 0.091 (95% CI 0.003-0.245)
-Mtb: EC50 = 0.320 (95% CI 0.155-0.604)
K48-linked poly-Ub (%maximum)
[MG-132] (μM)
d
+Mtb: EC50 = 8.337 μM (95% CI 5.577-12.46)
Mock: EC50 = 2.541 μM (95% CI 0.405-0.8471)
Cathepsin B activity (% of vehicle control)
[E64d] (μM)
e
DMSO
Bafilomycin
No lysotracker
Mock
+Mtb
10 μm
Lysotracker Red
GFP (Mtb)
f
Mock
+Mtb
Relative lysotracker+ area (% of DMSO control)
***
*
DMSO
Bafilomycin
DMSO
Bafilomycin

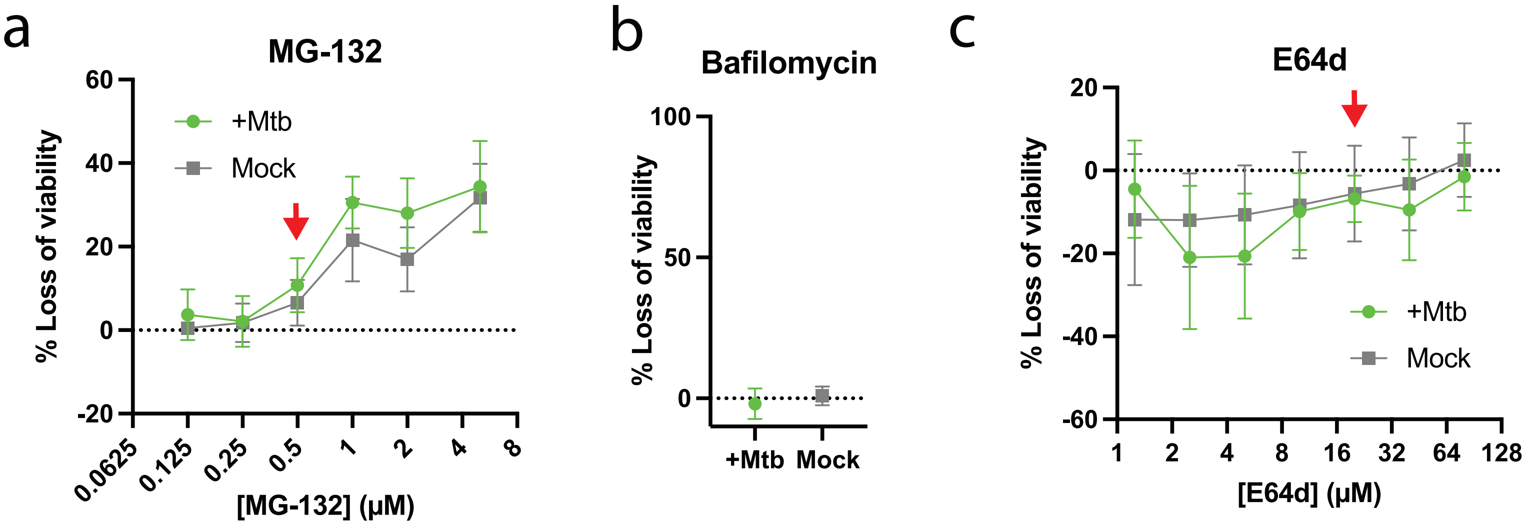

a
MG-132
% Loss of viability
60
40
20
0
-20
+Mtb
Mock
0.0625
0.125
0.25
0.5
1
2
4
8
[MG-132] (μM)
b
Bafilomycin
% Loss of viability
100
50
0
+Mtb
Mock
c
E64d
% Loss of viability
20
0
-20
-40
-60
+Mtb
Mock
1
2
4
8
16
32
64
128
[E64d] (μM)

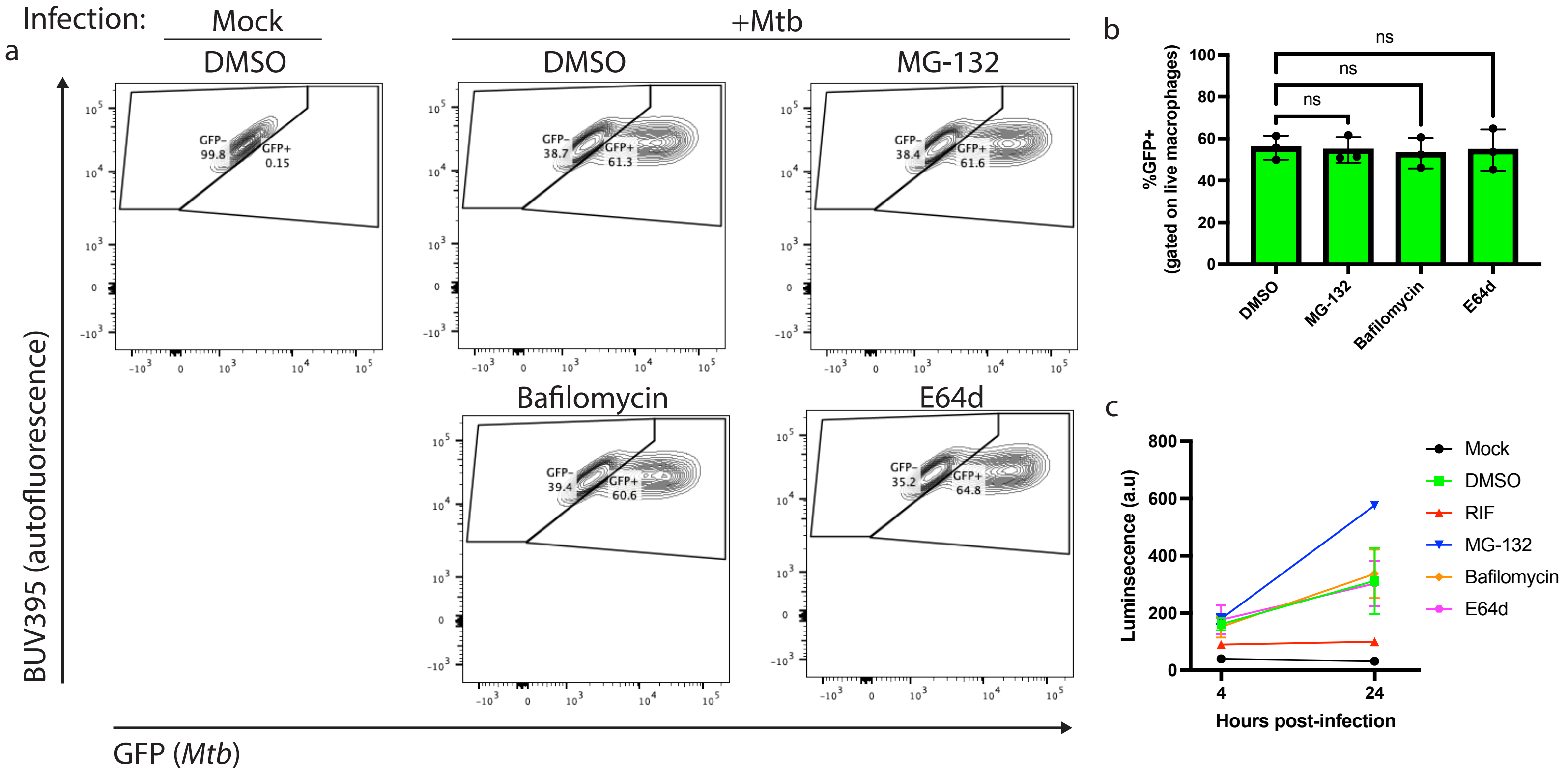
a
Infection:
Mock
+Mtb
DMSO
DMSO
MG-132
Bafilomycin
E64d
GFP-
99.8
GFP+
0.15
GFP-
38.7
GFP+
61.3
GFP-
38.4
GFP+
61.6
GFP-
39.4
GFP+
60.6
GFP-
35.2
GFP+
64.8
BUV395 (autofluorescence)
GFP (Mtb)
b
%GFP+
(gated on live macrophages)
ns
ns
ns
DMSO
MG-132
Bafilomycin
E64d
c
Luminsecence (a.u)
Hours post-infection
Mock
DMSO
RIF
MG-132
Bafilomycin
E64d
